# Prot-LAMBDA: Explicit Distance Learning Enhances Structural Reasoning in Protein Language Models

**DOI:** 10.64898/2026.08.23.746565

**Authors:** Nabil Ibtehaz, Zicong Zhang, Yuki Kagaya, Min Xu, Kentaro Tomii, Daisuke Kihara

**Affiliations:** Department of Computer Science, Purdue University, West Lafayette, IN 47906, USA; Department of Biological Sciences, Purdue University, West Lafayette, IN 47906, USA; Ray and Stephanie Lane Computational Biology Department, Carnegie Mellon University, PA 15213, USA; Artificial Intelligence Research Center (AIRC), National Institute of Advanced Industrial Science and Technology (AIST), 2-4-7 Aomi, Koto-Ku, Tokyo, 135-0064, Japan

## Abstract

Protein language models (PLMs) learn evolutionary information from large-scale sequence data, but three-dimensional relationships are encoded only implicitly. Here, we introduce Prot-LAMBDA (Protein LAnguage Model Boosted with Distance Awareness), a PLM that explicitly incorporates spatial relationships by coupling residue embeddings with inter-residue contacts. Prot-LAMBDA improves performance across diverse structure-related tasks, including contact, secondary structure, backbone geometry, solvent accessibility, and protein fold prediction. Notably, it achieves a twofold improvement in long-range contact recall and an 11.7% reduction in ψ-angle prediction error relative to ESM2-3B. Despite having approximately fivefold fewer parameters, Prot-LAMBDA also improves 3D structure prediction over ESM2-3B by 5–7% in TM-score when coupled to the same structure-prediction module. Building on these representations, we developed LambdaFold, a lightweight distance-guided structure prediction framework that achieves performance comparable to ESMFold on proteins strictly non-redundant to the training data. Finally, retrieval-augmented integration of structural templates increases mean TM-score substantially for targets with high template coverage and rescues several incorrect folds. Together, these results demonstrate that explicit spatial constraints enable efficient and generalizable structural representation learning and protein structure prediction.

## Introduction

Protein structure prediction has been studied for decades as one of the central problems in computational biology. Around the early 2000s, a variety of approaches were actively explored, including *ab initio* folding [1–3], template-based modeling [4–6], and ensemble approaches [7, 8] that combined multiple prediction strategies. During this period, evolutionary information derived from multiple sequence alignments (MSAs), particularly correlated mutation analysis, emerged as a promising direction for identifying residue-residue contacts in proteins [9, 10]. The idea of correlated mutations became especially powerful with the introduction of deep learning methods [11, 12]. Since then, the field has largely converged on a common framework: using deep neural networks to extract structural information from MSAs and predict residue contacts, inter-residue distances, and orientations [13, 14]. These advances culminated in highly successful systems such as AlphaFold2 (AF2) [15], which dramatically transformed structural biology and achieved near-experimental accuracy for many globular proteins.

Despite their remarkable success, MSA-based approaches have important limitations. Their performance strongly depends on the availability of sufficiently deep and diverse homologous sequence datasets [15]. Consequently, they often struggle on proteins with shallow MSAs, including orphan proteins [16], de novo designed proteins [17], and rapidly evolving viral proteins [18, 19]. Even extensive searches against large metagenomic databases may fail to identify enough homologous sequences for these targets. These limitations motivated the search for approaches that can infer structural information directly from amino acid sequences without relying heavily on evolutionary alignments.

Inspired by the success of large language models in natural language processing, protein language models (PLMs) have recently emerged as a promising alternative framework [20, 21]. Trained on millions of protein sequences using self-supervised learning objectives such as masked language modeling [22], PLMs learn statistical relationships, sequence patterns, and protein family-level semantics directly from sequence data [23, 24]. This intuition enabled the development of single-sequence structure prediction methods [25–27] such as ESMFold [28], which replace explicit MSA representations with embeddings generated by PLMs. Compared with MSA-based methods, PLM-based approaches are particularly advantageous for proteins with limited evolutionary information, including orphan and viral proteins.

However, most existing PLMs are fundamentally adapted from natural language modeling techniques and primarily learn sequence-level statistical patterns rather than explicit structural principles. Although proteins are inherently three-dimensional objects, conventional PLMs are generally trained without direct structural supervision and therefore do not possess an explicit understanding of protein geometry. Several recent studies have attempted to incorporate structural information into PLMs through fold-aware training [29], contrastive learning with inter-residue contact map similarity [30], structural tokenization [31], or coordinate-based objectives [32]. Nevertheless, these approaches remain relatively limited in their ability to directly encode spatial relationships and residue-level geometric constraints within the learned embedding space. Rather than modeling the local contextual relationships among constituent amino acids, these approaches primarily learn a comparative representation of a specific aspect of protein structures at a global level. The evaluation of such structural representation quality has also largely been restricted to a limited set of downstream tasks and primarily focused on global properties, such as protein fold prediction or function prediction. Most importantly, prior work has not been systematically investigated whether these embeddings are meaningful at a fine-grained residue level or whether they can support reasoning about the complete 3D structure of proteins.

Here, we investigate whether structural understanding can be directly extrapolated from a distance-regularized embedding space. We achieve this by correlating spatial distances with embedding differences, leading to the development of Prot-LAMBDA (Protein LAnguage Model Boosted with Distance Awareness). A key advantage of our structural representation learning framework over existing sequence-structure cross-modal constructive learning approaches is that, whereas prior methods typically learn implicit structural relationships through contrastive learning across proteins, our model encodes explicit pairwise distance information at the residue level. Incorporating this distance-based supervision during pretraining enables higher-order structural and geometric reasoning, as evidenced by consistent improvements in long-range contact prediction and enhanced modeling of dihedral angles. Furthermore, this local structural understanding naturally scales to global structural modeling, enabling accurate prediction of full, all-atom 3D protein structures. Our framework demonstrates strong versatility across diverse downstream structural tasks: For contact prediction, Prot-LAMBDA achieves consistent improvements on the most challenging distant residue contacts, outperforming both structure-aware PLMs and dedicated contact prediction methods by 6% and 3%, respectively. In structural understanding tasks, such as, secondary structure and fold prediction, our representations improve performance by 1–2% over standard PLMs without any task-specific fine-tuning. For geometry-oriented tasks, such as prediction of the dihedral psi angle, we observe a relative improvement of 11.74%. Finally, our embeddings support accurate end-to-end three-dimensional structure prediction, achieving gains of 5–7% compared with a PLM, ESM2-3B that is approximately five times larger.

Building on these structurally meaningful embeddings, we further developed LambdaFold, a protein 3D structure prediction framework that explicitly incorporates distance information and achieves performance comparable to ESMFold on challenging proteins despite using less than 10% of its parameters. Furthermore, we extended the method with a retrieval-augmented generation (RAG) system that iteratively updates predicted 3D structures by integrating distograms derived from available structural templates. This update is performed using a novel module that combines triangular cross-attention with distance-feedback driven invariant point attention (IPA) structure-modeling module. This implementation further improves the predicted models, enabling several proteins to adopt the correct fold and increasing the average TM-score by an additional 5%.

## Results

### Incorporating distance understanding into protein language model embedding

Prot-LAMBDA is based on the central premise that residue-level protein language model (PLM) embeddings encode latent three-dimensional relationships among amino acids. While existing structure-aware PLMs capture structural information implicitly, Prot-LAMBDA explicitly couples sequence representations with inter-residue spatial constraints, while retaining the biochemical and evolutionary information learned from large-scale protein sequence data. Instead of attempting to align embeddings with absolute Cartesian coordinates, which is ill-defined due to the lack of a canonical reference frame in protein structures, the model learns a relational representation in which embedding geometry reflects inter-residue spatial relationships. **Fig. 1a** shows network architecture.

**Fig. 1.**
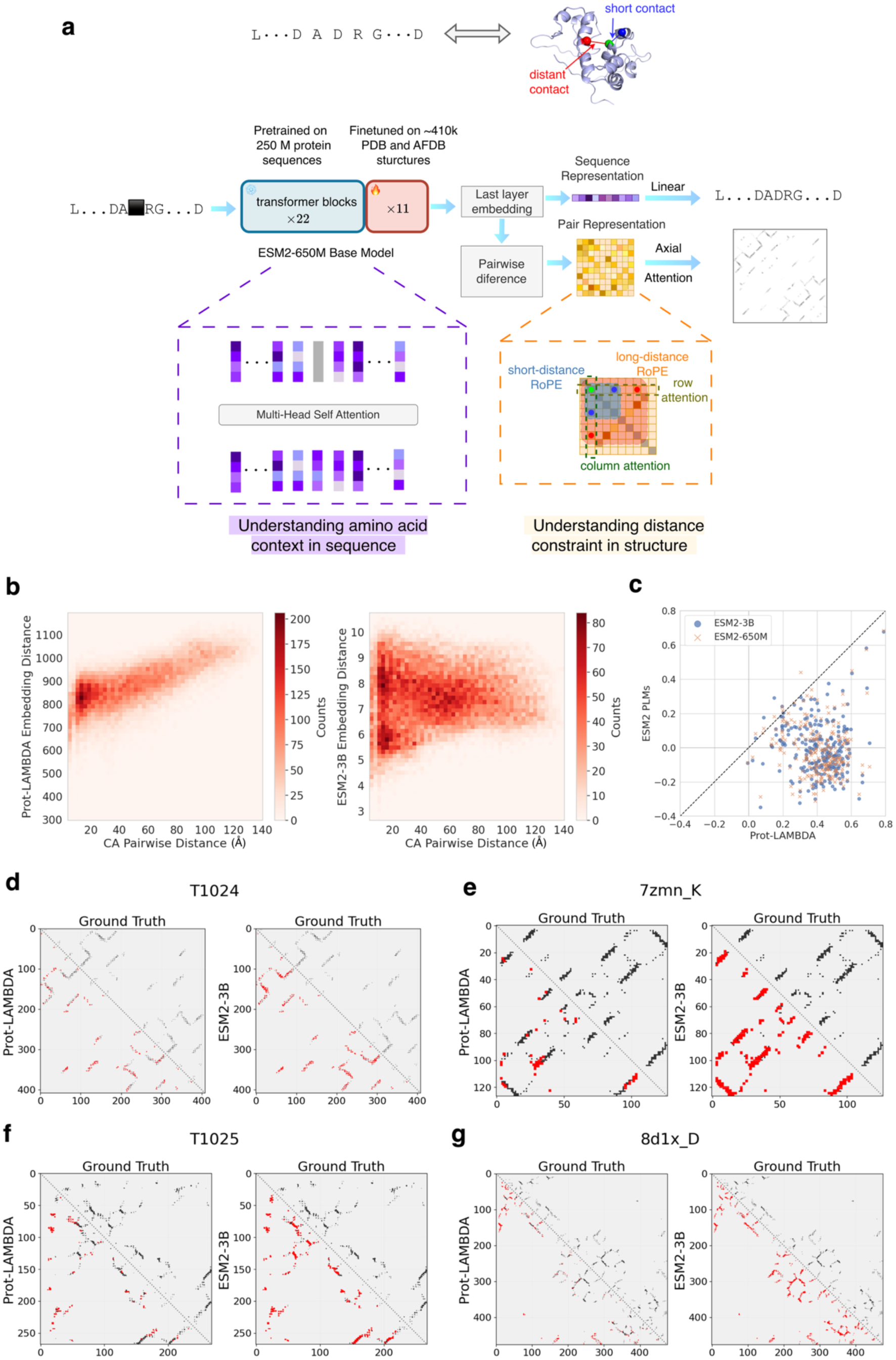
Prot-LAMBDA model, distance-aware representation and contact map prediction. **a.** Overview of the model architecture. Prot-LAMBDA bridges protein sequence context with structural distance constraints. The core architecture is a language model partially finetuned from a pretrained ESM2-650M model. The embedding from the final transformer layer serves as the sequence representation, while its pairwise difference forms the pair representation, which are utilized for masked language modeling and contact map prediction, respectively. The lower panel highlights key architectural components: standard multi-head self-attention progressively refines the sequence representation, whereas a single layer of axial attention, integrated with RoPE to account for sequence separation, refines the pair representation. **b–c.** Relationship between embedding differences and spatial distances. **b**. shows the correlation for the CASP14 target T1030 (spearman correlation of 0.55 for Prot-LAMBDA vs –0.1 for ESM2-3B). **c**. The summary across the CASP14 and CAMEO benchmark and comparison against ESM2 PLMs. Mean values: Prot-LAMBDA 0.41; ESM2-3B –0.006; ESM2-650M –0.014. **d-g.** Examples of predicted contact maps by Prot-LAMBDA and ESM2-3B. **d**. an α-class protein CASP14 target T1024 (LmrP PDB ID: 6t1z_A), length: 408 amino acids (aa), long precision (P) @ L scores (precision computed from the top L highest-probability predictions, where L is the protein length) 0.78 vs 0.69. **e**. a β-class protein, RECQL5 helicase (PDB ID: 7zmn_K), length: 127 aa, long P @ L scores 0.95 vs 0.54. **f**. an α/β-class protein, T1025 (AtmM, PDB ID: 6uv6_A), length: 268 aa, long P @ L scores 0.97 vs 0.79. **g**. an α/β-class protein, aminopeptidase A (PDB ID: 8d1x_D), length: 476 aa, long P @ L scores 0.99 vs 0.71. For each map, points above the diagonal line represent the ground-truth contacts, while points below the diagonal line correspond to the predicted contacts by Prot-LAMBDA (left) and ESM2-3B (right). Contacts missed in the predictions (false negatives) are highlighted in red. Contact prediction by ESM2-650M are provided in Supplementary Fig. 1.

### Sequence representation

We build upon a pretrained PLM that maps a protein sequence of length *L* into residue-level embeddings, producing a representation in ℝ^!×#^, where each row corresponds to an amino acid embedding. These models, trained via masked language modeling on millions of protein sequences (e.g., ESM-2; [28]), learn statistical regularities of sequence composition and implicitly capture aspects of protein structure. However, this structural information is not explicitly constrained. In Prot-LAMBDA, we fine-tuned ESM-2 (650M parameters), updating only the last 11 of 33 layers to balance efficiency and expressivity. The final-layer residue embeddings are extracted as the sequence representation, which encodes both local amino acid identity and global sequence context (**Fig. 1a**). We initially explored parameter-efficient fine-tuning strategies, including low-rank adaptation (LoRA) [33], but observed consistently inferior performance in this setting (data not shown).

### Pair representation and contact map learning

Our objective in the developing Prot-LAMBDA is to model the correspondence between embedding differences of residue pairs and their spatial separation in the three-dimensional space. To explicitly model structural relationships, we define a pair representation based on embedding differences between residues:

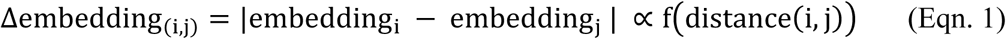

which is designed to correlate with inter-residue spatial proximity. Let *d_ij_* denote the Euclidean distance between residues *i* and *j* (defined using Cβ atoms, or Cα for glycine). Rather than regressing continuous distances, which proved unstable in preliminary experiments, we simplify the problem to binary contact prediction using:

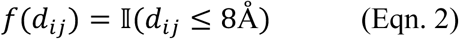

A naïve linear mapping from pairwise embedding differences to contacts is insufficient because residue interactions depend strongly on both local sequence separation and global structural context. To address this, we introduce a 2D interaction modeling module using axial attention [34], which factorizes full pairwise attention into row-wise and column-wise operations over the residue–residue matrix. For instance, as shown in Fig. 1a, the contact information between any residue pair (i, j) is inferred by separately attending along the row and column axes, effectively contextualizing the spatial constraints of the i^th^ and j^th^ residues against the entire protein. This enables efficient propagation of information across all residue pairs, allowing each interaction to be contextualized by the global contact landscape.

To encode sequence-order dependence, we incorporate rotary position embeddings (RoPE) [35] into the pair representation. RoPE applies position-dependent rotations to query and key vectors such that attention scores depend on relative sequence offsets. As a result, nearby residues experience small phase shifts (blue region, **Fig. 1a**), while distant residues undergo larger transformations (orange region, **Fig. 1a**), enabling explicit encoding of relative positional structure without additional learned parameters.

### Joint architecture and training

The sequence and pair representations are integrated within a unified architecture (**Fig. 1a**). The sequence branch is processed through multi-head self-attention (20 heads per layer), enabling progressive refinement of residue-level embeddings across 33 layers. The pair representation is then processed through axial attention (8 heads), enabling global contextualization of the contact map by jointly modeling row-wise and column-wise dependencies across all residue pairs.

The model is trained jointly using two objectives: masked language modeling (MLM) and contact map prediction. For MLM, 15% of residues are randomly masked and the loss is computed only on masked positions, following standard PLM training protocols [21]. To prevent degradation of pretrained sequence knowledge during contact-map training, i.e., catastrophic forgetting [36], we adopt a curriculum learning strategy. The contact loss is initially computed on 50% of residue pairs and gradually expanded to the full interaction set, stabilizing optimization and preserving pretrained representations.

Prot-LAMDA was trained on a training dataset, which combined experimentally determined protein structures from PDB [37] (**Supplementary Table S1**) with predicted structures from AlphaFold DB [38] (**Supplementary Table S2**), while rigorously excluding redundancy with 50 proteins from CASP14 [39] and 194 proteins from CAMEO [40] (from April 01 2022 through June 25 2022, as used in [15, 28]) benchmarks, two of the most widely used test sets to date. The PDB portion includes pre-May 2020 chains filtered by resolution, length, and sequence identity, with additional inclusion of NMR structures, resulting in ∼99k chains. The AFDB portion was derived from largest UniClust30 clusters (more than 30 member proteins), filtered based on size and confidence (pLDDT), yielding ∼310k proteins without further clustering. All sequences with more than 25% identity to benchmark datasets are removed to ensure a clean evaluation of generalization. Very importantly, unlike prior work, this approach explicitly enforces non-redundancy and uses a relatively smaller dataset to prioritize representation quality over memorization. During training, PDB and AFDB samples were mixed in a 1:2 ratio and processed using fixed-length crops of 256 residues (see Methods for more details). We used the same validation set as ESMFold, consisting of 776 CAMEO proteins released between August 2021 and January 2022 (**Supplementary Table S3**). The primary datasets used to evaluate our method are CASP14 and CAMEO (April 01 2022 to June 25 2022) for both contact map and structure prediction tasks. Additionally, we evaluated performance on structure understanding tasks using the TAPE [41], PEER [42] benchmarks.

Despite using a simpler formulation for incorporating distance awareness, the learned embeddings of Prot-LAMBDA seem to capture meaningful 3D spatial relationships. Specifically, in **Fig. 1b**, there is a clear positive correlation between Cα distance and embedding difference for residue pairs in an example protein T1030, indicating that spatial proximity is reflected in the representation. In contrast, embeddings from ESM2-3B show little such correlation. This trend generalizes across the CASP14 and CAMEO dataset, where the proposed embeddings achieve an average Spearman correlation of 0.41 (**Fig. 1c**), reflecting a moderate relationship between embedding space and physical distance. Meanwhile, ESM2 embeddings exhibit almost no correlation at all under the same evaluation (full data provided in **Supplementary Table S4**).

### Contact map prediction from single sequence information

We begin our evaluation by assessing contact map prediction performance, as contact map prediction serves as a pretraining objective of our model and provides a direct way to validate the effectiveness of the pretraining process. We evaluated our performance on the CASP14 and CAMEO (April 01, 2022 to June 25, 2022) benchmarks. As mentioned earlier, to ensure a rigorous assessment, we confirmed that there is no overlap with our training dataset and that the training data is non-redundant; specifically, no training sequence shares more than 25% similarity with the test proteins. We compared contact predictions by Prot-LAMBDA with ESM2-650 and ESM2-3B [28]. For the two ESM2 models, we used their officially released contact prediction head.

Four examples of contact prediction by Prot-LAMBDA and ESM2-3B are shown in **Fig. 1d-g**. Contacts that are missed in a prediction are highlighted in red. These proteins are from distinct structural topologies: an α-class protein (T1024; **Fig. 1d**), a β-class protein (7zmn_K; **Fig. 1e**), an α/β-class protein (T1025; **Fig. 1f**), and a large 477-residue protein (8d1x_D; **Fig. 1g**). Predictions with ESM2-650M for the same proteins are shown in **Supplementary Fig 1**. In the first example, ESM2-3B failed to predict the helical interactions between residues 54–66 and 111–121. In addition, it missed many residue-residue interactions involving secondary structure elements, resulting in extensive false negatives (red regions) across medium– and long-range contacts. Clearer differences were observed in **Fig. 1e–g**, where ESM2-3B missed most native interactions, in sharp contrast to the substantially more accurate predictions by Prot-LAMBDA. These qualitative differences were also reflected in the quantitative evaluation metrics (**Supplementary Fig. 1**). For instance, in terms of the most challenging long-range precision@L scores (contacts that are between residues more than 24 residues apart in the sequence are considered. Precision was computed for top L predictions with the highest probabilities, where L is the length of the protein), Prot-LAMBDA consistently outperformed the baseline method across all four targets. Specifically, Prot-LAMBDA achieved higher scores for T1024 (0.78 vs. 0.69), 7zmn_K (0.95 vs. 0.54), T1025 (0.97 vs. 0.79), and 8d1x (0.99 vs. 0.71). Other accuracy metrics are provided in **Supplementary Fig. 1**.

Overall performance on the CASP14 and CAMEO datasets is summarized in **Fig. 2a** and b, respectively. Individual data are provided in **Supplementary Table 1 and 2** and full raw data is provided in **Supplementary Table S5**. The radar plots show precision, recall, and F1 scores for short-, medium-, and long-range contacts, corresponding to residue pairs separated by 6–12, 12–24, and more than 24 residues in sequence, respectively. Following standard machine learning practice, predicted contact maps were binarized using a threshold of 0.5, and the resulting precision, recall, and F1 scores were computed against the ground-truth contact maps. As shown in **Fig. 2a** and **2b**, Prot-LAMBDA consistently outperforms both ESM2-650M and the much larger ESM2-3B across all metrics. The most pronounced gains are observed in recall and F1 score for medium and long-range contacts, which are among the most challenging to predict. Remarkably, Prot-LAMBDA achieves twice the recall of the ESM2 models (0.17-0.19 vs 0.39 for CASP14, **Fig. 2a**) and 0.24-0.27 vs 0.60 for CAMEO, **Fig. 2b**).

**Fig. 2.**
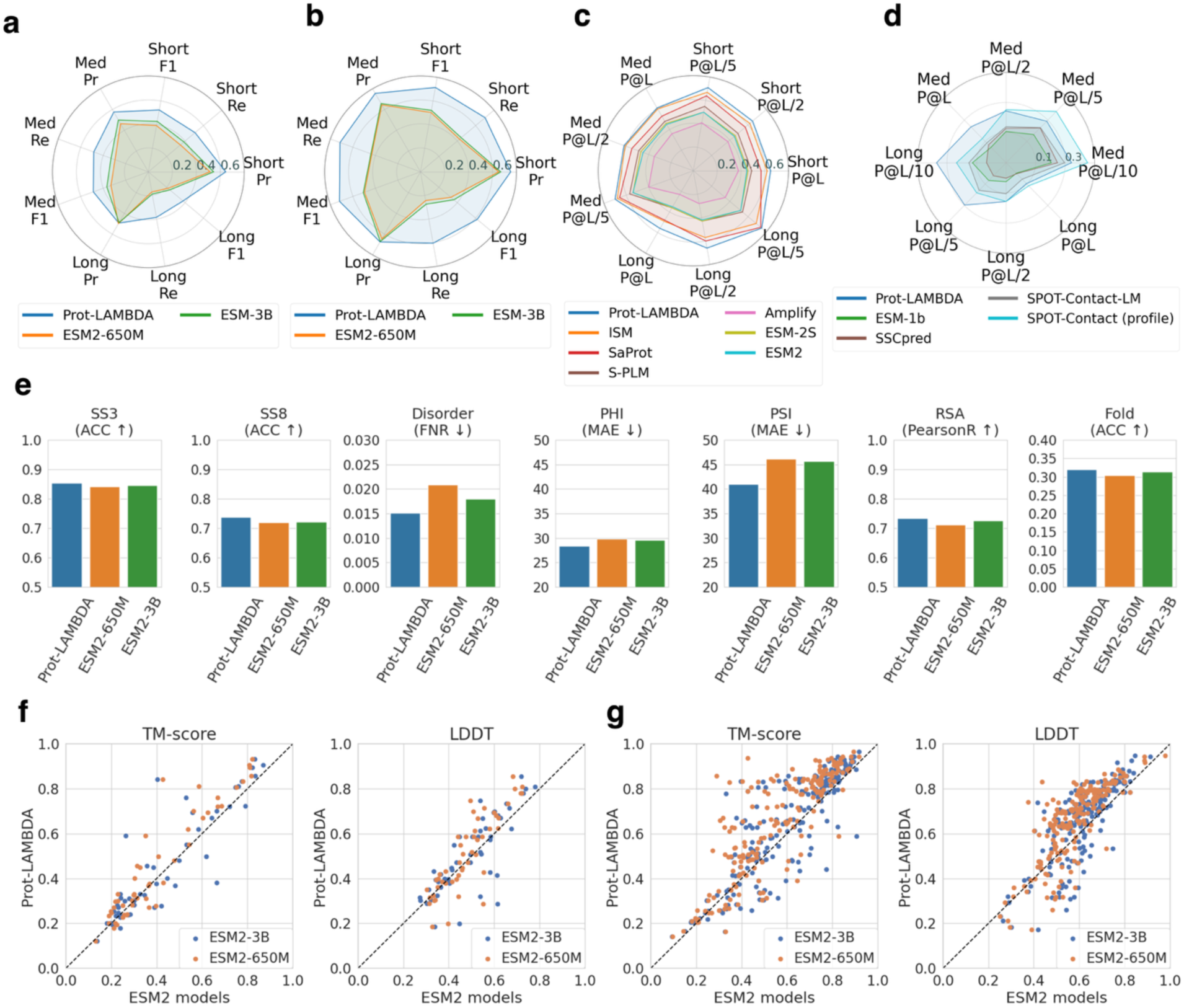
Structural understanding of the Prot-LAMBDA model. **a–d**. Contact map prediction performance. **a.** The CASP14 dataset (50 targets). Prot-LAMBDA short-, medium-, long-range precision (Pr), recall (Re) and F1 scores are compared with ESM2-650M and ESM2-3B models. Long range recall; Prot-LAMBDA, 0.39; ESM2-650M, 0.17; ESM2-3B, 0.19. More data in Supplementary table 1. **b**. The CAMEO dataset (194 targets). Prot-LAMBDA compared with ESM2-650M and ESM2-3B. Prot-LAMBDA achieves a long-range recall of 0.60 vs 0.24 (ESM2-650M) – 0.27 (ESM2-3B), More data in Supplementary Table 2. **c**. Comparison with 5 structure-aware PLMs along with ESM model on the TAPE benchmark with 40 proteins. Top L/k (k=1,2,5) contact map prediction precision scores across short-, medium– and long-ranges are reported. Prot-LAMBDA achieves 0.65, 0.61 and 0.56 for short, medium, long-range P@L, respectively. The second-best method ISM achieves 0.62, 0.6 and 0.49 for these 3 metrics, respectively. More data in Supplementary Table 3. **d**. Comparison on 15 CASP14-FM targets with dedicated single-sequence-based methods such as SPOT-Contact-LM, SSCPred. Prot-LAMBDA achieves long-range P@L/10 score of 35.35% vs. 18.92%, achieved by SPOT-Contact-LM, and surpasses profile-based SPOT-Contact on 5 of 8 metrics. More data in Supplementary Table 4. **e**. Prediction of 3-state (SS3) and 8-state (SS8) protein secondary structure, disordered region, PHI and PSI dihedral angles, relative solvent accessibility (RSA), and protein fold topology from the PEER benchmark. Comparison is against ESM2 models. Metrics include accuracy (ACC ↑) for 3-state (SS3) and 8-state (SS8) secondary structure and protein fold prediction; False Negative Rate (FNR ↓) for disorder; Mean Absolute Error (MAE ↓) for PHI and PSI backbone dihedral angles; and Pearson Correlation Coefficient (PearsonR ↑) for Relative Solvent Accessibility (RSA). Arrows indicate the direction of improved performance. More data in Supplementary Table 5. **f–g**. Evaluation of 3D structure prediction capability using frozen PLM embeddings coupled with an Invariant Point Attention (IPA) structure module. Scatter plots compare Prot-LAMBDA (y-axis) against ESM2-650M (orange crosses) and ESM2-3B (blue dots) on CASP14 (**f**) and CAMEO (**g**). Metrics shown are TM-score and LDDT for global and local structural accuracy, respectively. Average TM-score values: Prot-LAMBDA, 0.456; ESM2-650M, 0.408; ESM2-3B, 0.434 (CASP14) and Prot-LAMBDA, 0.657; ESM2-650M, 0.570; ESM2-3B, 0.614 (CAMEO). Average LDDT values: Prot-LAMBDA, 0.512; ESM2-650M, 0.483; ESM2-3B, 0.500 (CASP14) and Prot-LAMBDA, 0.662; ESM2-650M, 0.587; ESM2-3B, 0.618 (CAMEO).

Next in **Fig. 2c**, we compared it with four existing structure-aware PLMs. These four PLMs incorporated structure information in different aspects: ESM2-S involves protein fold topology in the pretraining [29], S-PLM compares protein sequences based on their contact map similarity [30], SaPort incorporates structural tokenization through 3di tokens [31], and ISM correlates sequence embeddings with 3D coordinates [32]. To assess our model in this context, we evaluated contact map prediction performance using the TAPE benchmark [41], which is a general assessment benchmark for PLMs and their contact map evaluation framework has been adopted by these PLMs [31, 32]. Following the benchmark protocol, we trained Prot-LAMBDA on the provided training split by freezing the entire PLM and fine-tuning only the final linear layer and then evaluated the model on the corresponding test split [41], consistent with the training procedure used by the compared methods [31, 32]. Predictions were evaluated with short-, medium-, and long-range Precision@L/k (L is the length of protein and k=1, 2, 5) scores following the prior works [31, 32], (**Fig. 2c**; individual data in Supplementary Table 3). Prot-LAMBDA consistently outperforms all these structure-aware PLMs across all metrics. The results are particularly noteworthy because Prot-LAMBDA outperforms both SaProt, which uses explicit 3di [43] structural tokens as input, and ISM, which was structure-tuned on over five million structures.

In **Fig. 2d**, we further compared Prot-LAMBDA against a dedicated, state-of-the-art method SPOT-Contact-LM [44] (data in **Supplementary Table 4**). SPOT-Contact-LM employs a similarly sized PLM, incorporates additional structural features from SPOT-1D-Single, and uses a 12-layer deep residual network for contact map prediction. We used the same experimental protocol of SPOT-Contact-LM [44]. For a fair comparison, we finetuned Prot-LAMBDA on their training dataset [45] (similar to the previous experiment (**Fig. 2c**), only the topmost linear layer was modified, and rest of the model was frozen). Evaluation was performed on the same 15 targets from the CASP14-FM set using the same metrics used in the SPOT-Contact-LM work. We also included two competing methods considered by SPOT-Contact-LM, namely, single-sequence-based SSCPred [46], ESM-1b [21], and a profile-based variant of SPOT-Contact. We computed the same metrics used in the SPOT-Contact-LM paper [44]. Prot-LAMBDA consistently outperforms SPOT-Contact-LM across all metrics, achieving an impressive twofold improvement on long-range P@L/10 (35.35% vs. 18.92%). Moreover, Prot-LAMBDA is competitive with the profile-based SPOT-Contact, surpassing or matching it on five of eight metrics (2/4 for medium-range and 3/4 for long-range).

### Predicting other structural features and the tertiary structure of proteins

Subsequently in **Fig. 2e**, we performed other structure-related prediction tasks with Prot-LAMBDA. We used the PEER [42] benchmark, which provides training and testing datasets for predicting secondary structure (both 3 and 8 states), disorder, Phi, Psi dihedral angles, and Relative Solvent Accessibility (RSA). For all tasks, we perform linear probing (training a linear layer) on embeddings from Prot-LAMBDA and the baseline ESM2 models, using either globally pooled representations (for fold prediction) or residue-level embeddings (for the remaining tasks). Training follows the train–validation splits provided by PEER (see Methods).

First, we observe that model scale improves structural understanding in protein language models, as the larger ESM2-3B consistently outperforms the smaller ESM2-650M across all tasks. Nevertheless, Prot-LAMBDA embeddings capture even stronger structural information, consistently outperforming the larger ESM2-3B model by a modest margin. The largest improvement is observed for ψ-angle prediction, where Prot-LAMBDA reduces the mean absolute error by 11.74% relative to ESM2-3B. These gains are particularly noteworthy because Prot-LAMBDA was not explicitly trained on any of these downstream prediction tasks.

The results in **Fig. 2e** suggest that aligning and regularizing the embedding space with distance-based constraints implicitly encodes structural relationships, yielding protein representations with enhanced structural information.

To further assess the structural information encoded by Prot-LAMBDA embeddings, we evaluated their ability to support *de novo* protein structure prediction using a simple AF2-based architecture. Specifically, we attached a pretrained Invariant Point Attention (IPA) structure module [15] to frozen PLM embeddings and trained only the bridging layers and structure module (**Supplementary Fig. 2**). ESM2-650M and ESM2-3B, the latter serving as the backbone of ESMFold, were used as baselines. This setup can be viewed as a probing experiment to evaluate how well PLM embeddings align with AF2’s structural representation. All models were trained on the same set of approximately 98,000 PDB structures using the standard AlphaFold loss (excluding the violation loss; see Methods) and evaluated on the CASP14 and CAMEO test sets. Performance was assessed using TM-score [47], which measures global structural similarity, and LDDT (Local Distance Difference Test), which evaluates local structural accuracy [48].

As shown in **Fig. 2f** and **2g**, Prot-LAMBDA embeddings consistently improve structure prediction performance on both the CASP14 and CAMEO datasets over the ESM models. Compared with the base ESM2-650M model, Prot-LAMBDA achieves relative improvements of 11.7% in TM-score and 6.0% in LDDT on CASP14, and 15.3% and 12.8%, respectively, on CAMEO, demonstrating the benefit of embedding alignment with distance-based structural information. Notably, Prot-LAMBDA also outperforms the approximately 5-times larger ESM2-3B model, improving TM-score by 5.0% and LDDT by 2.4% on CASP14, and by 7.0% and 7.1%, respectively, on CAMEO (individual data in **Supplementary Table S6**).

### Extending the language model to 3D structure prediction

Building upon the strong structural understanding of Prot-LAMBDA, we developed LambdaFold, a lightweight protein structure prediction pipeline grounded in explicit distance modeling (**Fig. 3a**). Our design emphasizes efficiency and structured reasoning over protein conformational space, rather than memorizing structural patterns through extremely large model architectures. Like AF2, LambdaFold consists of a folding trunk followed by a structure module. However, unlike existing PLM-based methods such as ESMFold [28] and HelixFold [25], whose folding trunks rely primarily on implicit structural supervision, LambdaFold is explicitly designed to model residue-residue distance relationships throughout the folding process.

**Fig. 3.**
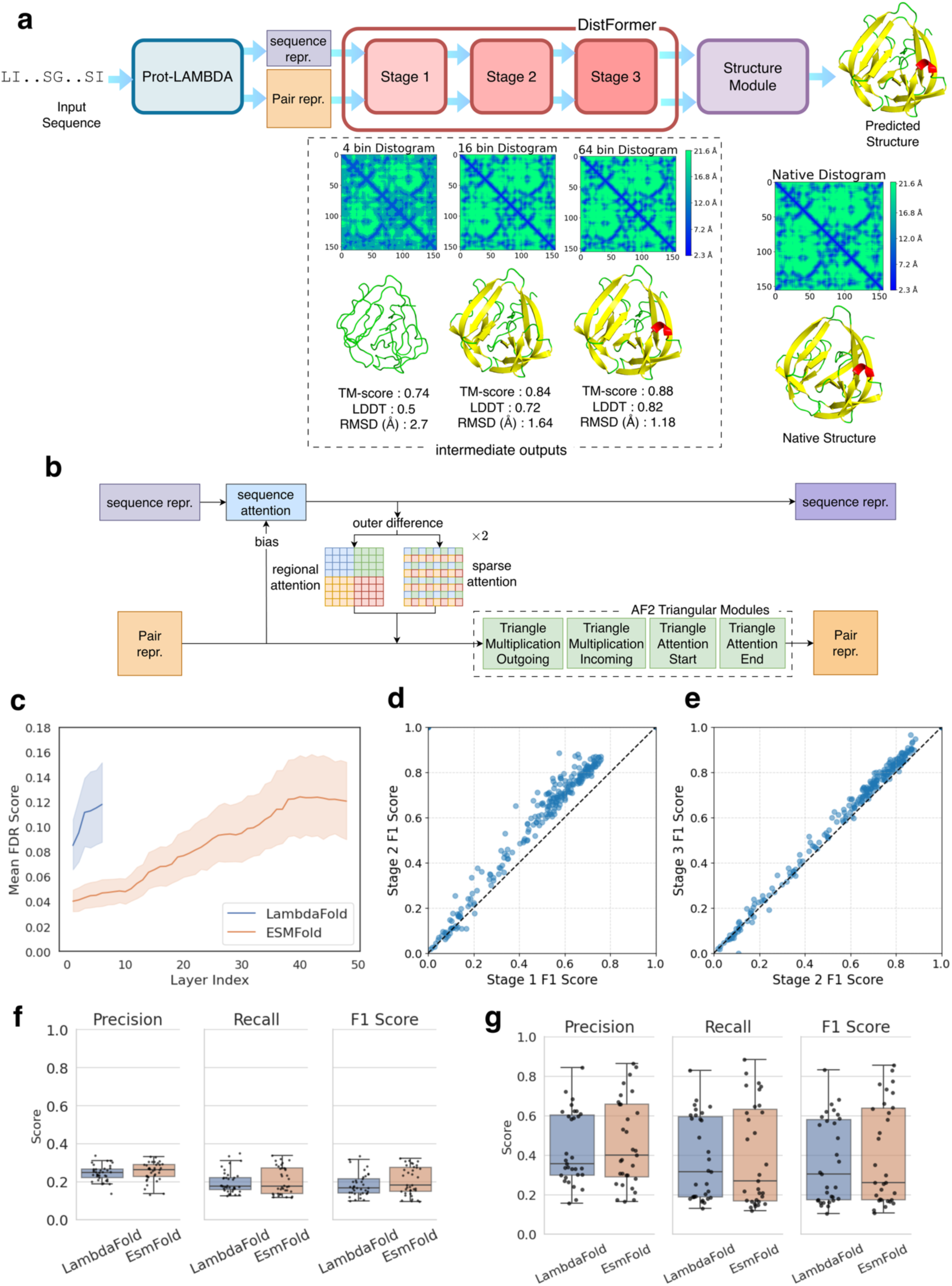
The LambdaFold model. **a**. Sequence and pair representations from Prot-LAMBDA are processed by DistFormer, a compact distance-guided folding trunk with three stages of progressive refinement, generating distograms (4, 16, and 64 bins) and corresponding 3D structures. An example from CASP target T1034 illustrates increasing structural accuracy across stages, with improvements in TM-score, LDDT, and RMSD. The progressive development and refinement of secondary structures in successive stages can be observed. Structures are colored by secondary structure classes: red for α helices, yellow for α sheets, and green for other regions.) **b**. Detailed architecture of a single DistFormer layer. Sequence representations are updated through attention biased by pair representations. Pairwise features are updated using regional and sparse attention to capture intra– and inter-domain geometric information, followed by AF2-style triangular updates to maintain geometric consistency. **c**. Efficacy analysis showing Mean False Discovery Rate (FDR) score relative to the layer index. Blue, LambdaFold (three layers); red, ESMFold (48 layers). **d–e.** Gradual improvement of contact formation in predicted 3D structures. The F1 score is computed for structures predicted at different stages against the ground truth. D and E present comparative views of correct contact predictions for stage 1 (mean F1 score 0.48) vs. stage 2 (mean F1 score 0.59) and stage 2 vs. stage 3 (mean F1 score 0.62), respectively. **f-g**. Comparison of LambdaFold and ESMFold on distogram prediction. **f**. the comparative evaluation of the two methods on 37 FM/TBM domains from the CASP14 dataset. (Mean values for Precision, Recall, and F1 score are 0.24, 0.20, and 0.18 for LambdaFold, vs. 0.25, 0.21, and 0.20 for ESMFold. Median values for Precision, Recall, and F1 score are 0.25, 0.18, and 0.17 for LambdaFold, vs. 0.26, 0.17, and 0.18 for ESMFold.), **g**. distogram comparison on 30 non-redundant proteins from the CASP14 and CAMEO datasets with respect to the ESMFold training set. The mean Precision, Recall, and F1 score: 0.44 vs. 0.46, 0.38 vs. 0.39, and 0.37 vs. 0.40 for LambdaFold and ESMFold, respectively. The median Precision, Recall, and F1 score: 0.35 vs. 0.39, 0.29 vs. 0.25, and 0.27 vs. 0.26 for LambdaFold and ESMFold, respectively.

To improve structural fidelity, we hierarchically decomposed distance prediction from coarse to fine resolution. We developed DistFormer (**Fig. 3a, 3b**), a compact six-layer architecture (compared with 48 layers in ESMFold) organized into three sequential two-layer blocks. The blocks progressively refine inter-residue distance (between the range 2.3–21.6 Å) predictions from 4 to 16 to 64 bins, with the final resolution matching that of ESMFold and AF2.

The design and training of DistFormer (**Fig. 3b**) introduce several modifications over conventional folding trunks [15, 28]. We retain the AF2-style framework of sequence–pair representation updates and triangular updates to maintain geometric consistency. However, instead of relying solely on elementwise integration of sequence and pair features, we incorporate regional and sparse attention inspired by vision transformers [49]. These mechanisms explicitly model local and long-range residue interactions, corresponding to intra-domain and inter-domain distance relationships, respectively. We hypothesize that this distance-aware design enables efficient structural reasoning with a lightweight architecture, reducing the need for the deep networks used in existing methods. Regional attention operates on local windows to capture intra-domain geometric patterns, whereas sparse attention uses a global grid-like pattern to model long-range inter-domain interactions [50]. We hypothesize that explicitly encoding both spatial scales enables more effective distance-aware representations with shallow architecture. Although these attention mechanisms increase parameters by ∼18.7%, DistFormer reduces the overall folding trunk size from 686M (ESMFold) to 58M parameters (∼12-fold reduction) by using only 6 layers instead of 48. The resulting distance representations are further refined through AF2-style triangular modules.

Furthermore, DistFormer was trained with explicit distance supervision. Pair representations from the final layer of each block (layers 2, 4, and 6) predict inter-residue distances (2.3–21.6 Å) at progressively finer resolutions of 4, 16, and 64 bins. To support this hierarchical refinement, the sequence and pair embedding dimensions are gradually increased across blocks (sequence: 512, 640, 768; pair: 192, 224, 256). In addition to distance prediction, full-atom structures are generated using a shared-weight AF2-style structure module applied after each block. The three blocks are trained sequentially: the first block is trained with an AF2-style loss, combining the standard AF2 3D structure prediction loss (FAPE loss, angle loss, violation loss) with our modified distogram prediction loss (product of huber and cross-entropy loss, see Methods), followed by the second and third blocks with earlier blocks frozen (see Methods). This staged design enables a modular architecture with flexible performance–complexity trade-offs.

To assess the quality of the learned pair representations across DistFormer layers, we analyzed their separability with respect to inter-residue distance bins using the Fisher Discriminant Ratio (FDR) [51] (see Methods). FDR quantifies feature discriminability as the ratio of inter-class separation to intra-class variability. To ensure a fair comparison between models with different feature dimensionalities, pair representations were first projected to a common latent space using Linear Discriminant Analysis (LDA) [52]. Fig. 3c shows the layer-wise FDR values averaged over 37 challenging FM/TBM domains from CASP14. Higher FDR values indicate better separation of distance classes and thus more informative representations for distance prediction. DistFormer begins with a substantially higher FDR (>0.08) than ESMFold (∼0.04), reflecting the distance-aware pretraining of Prot-LAMBDA compared to the generic masked language modeling objective used in conventional PLMs. Furthermore, the FDR increases more rapidly across DistFormer layers, indicating that structural information is accumulated more efficiently. Although ESMFold eventually reaches a slightly higher final FDR, it requires approximately eight times more layers and substantially more training data to achieve comparable representation quality. These results demonstrate that the distance-guided architecture and pretraining strategy of DistFormer produce highly discriminative structural representations with considerably greater efficiency.

The progressive improvement in representation quality also translates into more accurate structure prediction. To quantify this, we evaluated the contact maps derived from the predicted 3D structures at each of the three DistFormer stages using the CASP14 and CAMEO benchmark proteins (**Fig. 3d, 3e**) (invidual data in **Supplementary Table S7**). Contacts were defined using an 8 Å Cβ–Cβ distance threshold (Cα for glycine) and compared with the native contact maps using the F1 score. As shown in **Fig. 3d–e**, contact accuracy improves steadily across stages, with the mean F1 score increasing from 0.48 at Stage 1 to 0.59 at Stage 2 and 0.62 at Stage 3. These results demonstrate that the progressive refinement of the internal representations directly translates into increasingly accurate three-dimensional structures.

### Evaluation of distance prediction

**Fig. 3f** evaluates the accuracy of inter-residue distance prediction. Following the CASP14 assessment protocol [53], we converted the predicted distograms, consisting of 64 distance bins spanning 2.3–21.6 Å, into 2 Å bins covering the range of 4–20 Å. A residue pair was considered correctly predicted if the most probable distance bin matched the corresponding bin in the native structure. Residue pairs separated by fewer than six positions in sequence were excluded, as these short-range distances are relatively trivial to predict. Precision, recall, and F1 scores were computed for each target using a one-vs-all strategy for the multiclass distance-bin prediction. For each distance bin, residue pairs belonging to that bin were treated as positives and all other pairs as negatives. Thus, a correct prediction of the target bin was counted as a true positive (TP), prediction of another bin for a pair belonging to the target bin as a false negative (FN), and prediction of the target bin for a pair belonging to another bin as a false positive (FP); all remaining predictions were true negatives (TN). Precision, recall, and F1 were calculated separately for each distance bin and then averaged across bins to obtain the macro-averaged scores. Consistent with the CASP14 assessment and previous studies [53, 54], the evaluation was performed on the 37 FM and FM/TBM domains to focus on the most challenging prediction targets.

LambdaFold and ESMFold (without recycle) achieved comparable mean precision and recall in inter-residue distance prediction, with precision of 0.24 and 0.25, and recall of 0.20 and 0.20, respectively. Although ESMFold attained a somewhat higher mean F1 score (0.20 vs. 0.18 for LambdaFold), the performance gap was modest. Median values of these metrics by the two methods are also very close. Considering that LambdaFold employs a substantially smaller model, these comparable results are particularly noteworthy, demonstrating that its distance-aware representations enable competitive distance prediction despite its much lower model complexity (full data is provided in **Supplementary Table S8**).

We further evaluated distogram prediction on a more stringent benchmark comprising 30 CASP14 and CAMEO targets sharing less than 25% sequence identity with the ESMFold training set (**Fig. 3g**). On this non-redundant dataset, the performance gap between the two methods became even smaller. LambdaFold and ESMFold achieved nearly identical mean precision (0.44 vs. 0.46), recall (0.38 vs. 0.39), and F1 score (0.37 vs. 0.40). Despite slightly lower mean scores, LambdaFold attained higher median recall (0.29 vs. 0.25) and median F1 score (0.27 vs. 0.26), indicating more consistent performance across targets (individual data is provided in **Supplementary Table S9**).

These results suggest that the performance advantage of ESMFold is reduced when evaluated on proteins that are explicitly non-redundant with its training data, raising the question of how much of its performance derives from true generalization versus familiarity with large-scale training datasets. Nevertheless, considering that LambdaFold is substantially smaller and trained on a much smaller, carefully curated non-redundant dataset, its comparable performance demonstrates the effectiveness of its distance-guided representation learning.

### 3D structure prediction by LambdaFold

We next evaluated the structure prediction performance of LambdaFold. As illustrated in **Fig. 3a**, a three-dimensional structure is generated at each DistFormer stage, enabling progressive refinement of the predicted model. The prediction accuracy at the three stages is summarized in **Fig. 4a**. For comparison, we also included the results obtained by fine-tuning a pretrained AF2 structure module using only Prot-LAMBDA embeddings (**Fig. 2f–g**), allowing us to isolate the contribution of the DistFormer module beyond the structure-aware PLM.

**Fig. 4.**
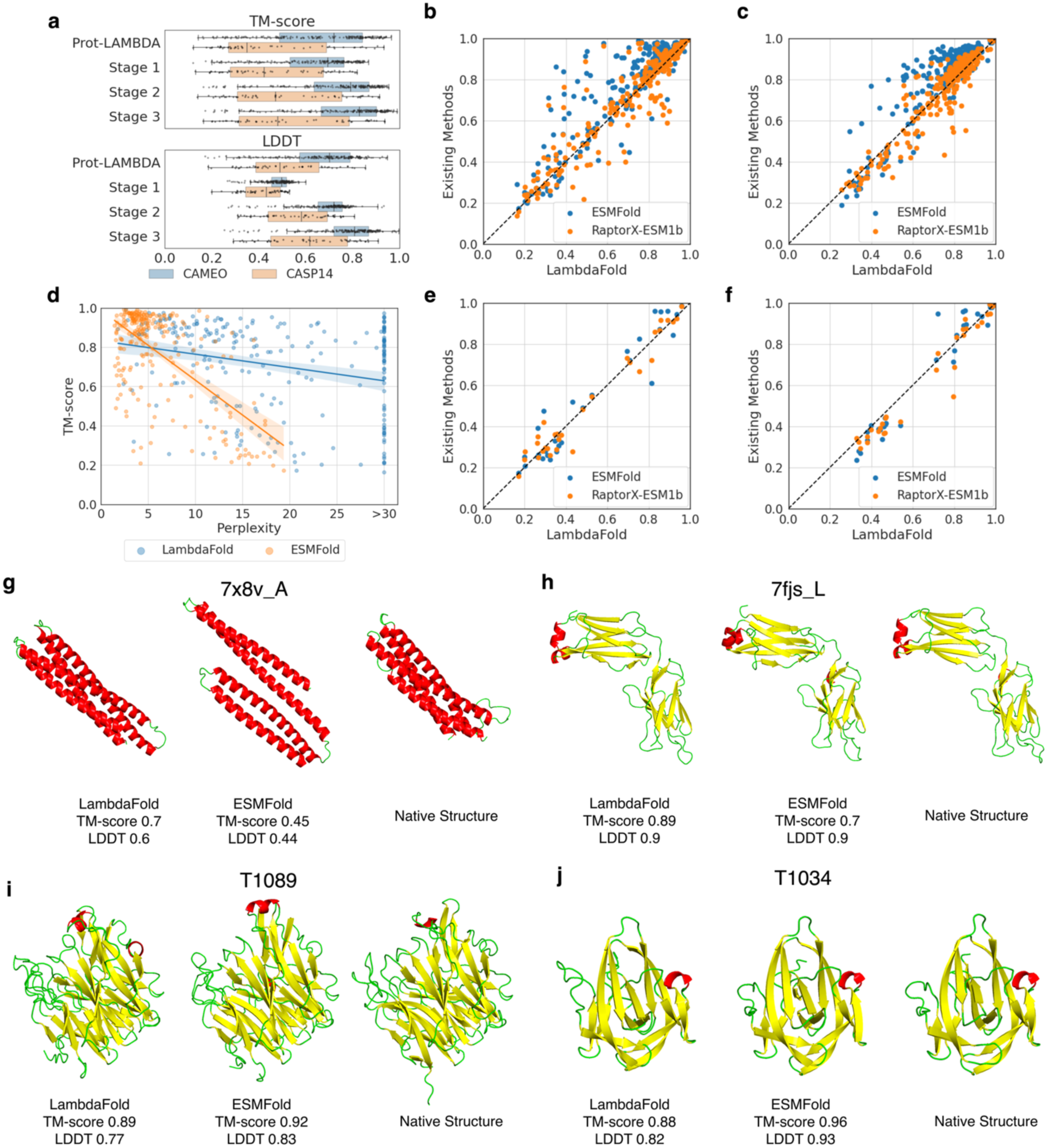
Protein 3D structure prediction by LambdaFold. **a.** Distribution of TM-scores (top) and LDDT (bottom) for three DistFormer stages and an AF2 structure module coupled with Prot-LAMBDA embedding. Two datasets, 50 targets of CASP14 (blue) and 194 targets of CAMEO (orange), were used. Boxplots indicate the median (center line), interquartile range (box), and 1.5x interquartile range (whiskers). **b-c**. Comparison on structure prediction performance with ESMFold (blue) and RaptorX-ESM1b (orange) on the combined 244 CASP14 and CAMEO benchmark proteins. **b**. TM-score (Average: LambdaFold, 0.71; RaptorX-ESM1b, 0.72; ESMFold, 0.78). **c**. LDDT (Average: LambdaFold, 0.74, RaptorX-ESM1b, 0.73; ESMFold, 0.81). **d**. Relationship between TM-score and sequence perplexity for LambdaFold (blue) and ESMFold (orange) for the combined 244 CASP14 and CAMEO targets. **e-f.** Head-to-head comparison of TM-scores (**e**) and LDDT (**f**) between LambdaFold and existing methods (ESMFold, blue; RaptorX-ESM1b, orange) on a non-redundant subset of 30 proteins from CASP14 and CAMEO benchmarks. **e**. TM-score. Average: LambdaFold, 0.51; RaptorX-ESM1b, 0.51; ESMFold, 0.52. **f**. LDDT. Average: LambdaFold, 0.663; RaptorX-ESM1b, 0.63; ESMFold, 0.65. **g**. predicted structures for 7×8v_A. Models of LambdaFold, EMSFold and the native structure are shown. TM-score and LDDT are shown below each model. **h**. predicted structure for 7fjs_L. **i**. T1089 CASP target. **j**. T1034 CASP target.

For both TM-score and LDDT, LambdaFold Stage 1 already surpassed the Prot-LAMBDA baseline, indicating that a single DistFormer stage effectively transforms the learned representations into more accurate structures. Additional DistFormer stages further refined the predictions, resulting in consistent improvements in both metrics. For example, in terms of TM-score, the average TM-score and LDDT increased by 8.27% and 19.48%, respectively, from Stage 1 to Stage 2, followed by further gains of 2.52% and 7.81% from Stage 2 to Stage 3. Overall, the average TM-score on the CASP14 dataset improved from 0.469 to 0.543, with the average LDDT increasing from 0.414 to 0.611. Similar improvements were observed on the CAMEO dataset, where TM-score increased from 0.33 to 0.749 and LDDT from 0.478 to 0.771 (individual data in **Supplementary Table S10**).

In **Fig. 4b–c**, we compared LambdaFold with two existing single-sequence structure prediction methods, ESMFold [28] and RaptorX-ESM1b [26], which uses a similarly sized ESM-1b backbone. Notably, the folding trunks of RaptorX-ESM1b and ESMFold are approximately four– and eight-fold deeper than that of LambdaFold, respectively. Despite its shallower architecture, LambdaFold closely matched RaptorX-ESM1b, achieving a mean TM-score of 0.71 versus 0.72 and a slightly higher mean LDDT of 0.74 versus 0.73. ESMFold achieved the highest performance, with mean TM-score and LDDT values of 0.78 and 0.81, respectively (invididual data in **Supplementary Table S11**). All these results were generated without any recycling, as our shallow Distformer is further disadvantaged by the much deeper networks.

As noted above, another important difference between LambdaFold and ESMFold/RaptorX-ESM1b is the composition of their training datasets. LambdaFold was trained under strict non-redundancy constraints, excluding proteins sharing more than 25% sequence identity with the benchmark targets. In contrast, ESMFold and RaptorX-ESM1b did not consider explicit sequence identity of other non-redundancy cutoffs. Indeed, we found that nearly 87% of the CASP14 and CAMEO test proteins share at least 25% sequence identity with proteins in the ESMFold training set.

To examine how training-set familiarity affects prediction accuracy, we analyzed TM-score as a function of perplexity (**Fig. 4d**). Perplexity reflects how well a PLM represents a given sequence, with lower values generally indicating greater familiarity with its learned sequence distribution. Approximately 84% of the test proteins had perplexity below 10 for ESMFold, compared with only 23% for Prot-LAMBDA, indicating that the benchmark sequences are substantially less familiar to our model. ESMFold showed a stronger dependence of prediction accuracy on perplexity: TM-score correlated with perplexity at −0.63 for ESMFold compared with −0.27 for LambdaFold (individual data in **Supplementary Table S12**). ESMFold performance declined markedly above a perplexity of 10, whereas LambdaFold maintained relatively stable performance up to approximately 20. These results suggest that LambdaFold is less sensitive to sequence familiarity and generalizes more robustly to sequences distant from its training distribution. The competitive structure prediction accuracy achieved under these conditions supports the effectiveness of distance-aware representation learning for generalizable protein structure prediction.

The above hypothesis is further supported by the prediction performance on a subset of 30 non-redundant proteins from the combined CASP14 and CAMEO datasets (**Fig. 4e, f**). On this strictly non-redundant subset, the performance gap narrowed considerably. LambdaFold achieved a comparable average TM-score (0.512) to RaptorX-Single (0.512), and the gap relative to ESMFold (0.521) was also reduced. In terms of LDDT, LambdaFold achieved an even higher average score (0.66) than both RaptorX-ESM1b (0.63) and ESMFold (0.65) (individual data in **Supplementary Table S13**).

In the next four panels (**Fig. 4g–j**), we show examples of models predicted by LambdaFold in comparison with those predicted by ESMFold. The first two cases (**Fig. 4g, 4h**) are examples in which LambdaFold performed better than ESMFold. The example shown in **Fig. 4g** is an α-helical protein (PDB ID: 7×8v_A). LambdaFold correctly modeled the tightly packed α-helical bundle, whereas ESMFold failed to properly pack the helices, resulting in a substantial difference in TM-score (0.70 vs. 0.45). In **Fig. 4h**, we show an antibody structure (PDB ID: 7fjs_L). In the LambdaFold prediction, the β-sheet formation in the β-sandwich at the top closely follows the native structure. In contrast, the ESMFold prediction deviates from the native β-sheet arrangement and also incorrectly forms α-helices, resulting in a lower TM-score of 0.70 compared with 0.89 for LambdaFold.

The next two examples (**Fig. 4i, j**) illustrate common cases in which LambdaFold achieved slightly lower TM-scores than ESMFold. For target T1089 (**Fig. 4i**), the two models performed comparably (TM-score: 0.89 vs. 0.92). LambdaFold accurately captures the core β-sheet architecture, with deviations mainly in flexible coil regions. This reflects a common pattern: LambdaFold reliably reconstructs structured cores but is less precise in flexible regions, where multiple conformations may be structurally plausible. A similar trend is observed for target T1034 (**Fig. 4j**). Both models capture the structured core but differ in flexible regions, where ESMFold more closely matches the native structure, resulting in higher scores (TM-score: 0.96 vs. 0.88; LDDT: 0.93 vs. 0.82). These examples suggest that the closer agreement of ESMFold in flexible regions may partly benefit from structural patterns memorized from its extensive training data, although further investigation is needed to understand this effect.

### Retrieval Augmented Generation (RAG) using templates

A primary limitation of our current model is its relatively small training dataset, which reduces performance on sequences distant from the training distribution [55]. To address this limitation, we explored retrieval-augmented generation (RAG), which supplements model predictions with relevant information retrieved from external databases [56, 57]. Similar to AF2, we developed a template search as a RAG strategy tailored to LambdaFold.

An overview of the LambdaFold-RAG pipeline is shown in **Fig. 5a**. Following the AF2 template-search strategy, structural templates are retrieved from PDB70 using MMseqs2 [58] and refined with HHsearch [59] through the ColabFold pipeline [60] with strict temporal cutoffs to prevent data leakage (see Methods). Retrieved templates are converted into binned pairwise Cβ distograms similar to AF2 (3.25–50.75 Å; 39 bins). Because alignment gaps leave portions of the template distogram undefined, we developed a template fusion module that integrates available template information with the distogram obtained from LambdaFold predicted structure. The fused distogram is then used to guide a modified invariant point attention (IPA) module, which first optimizes Cβ positions according to the distance constraints and subsequently refines all remaining atoms.

**Fig. 5.**
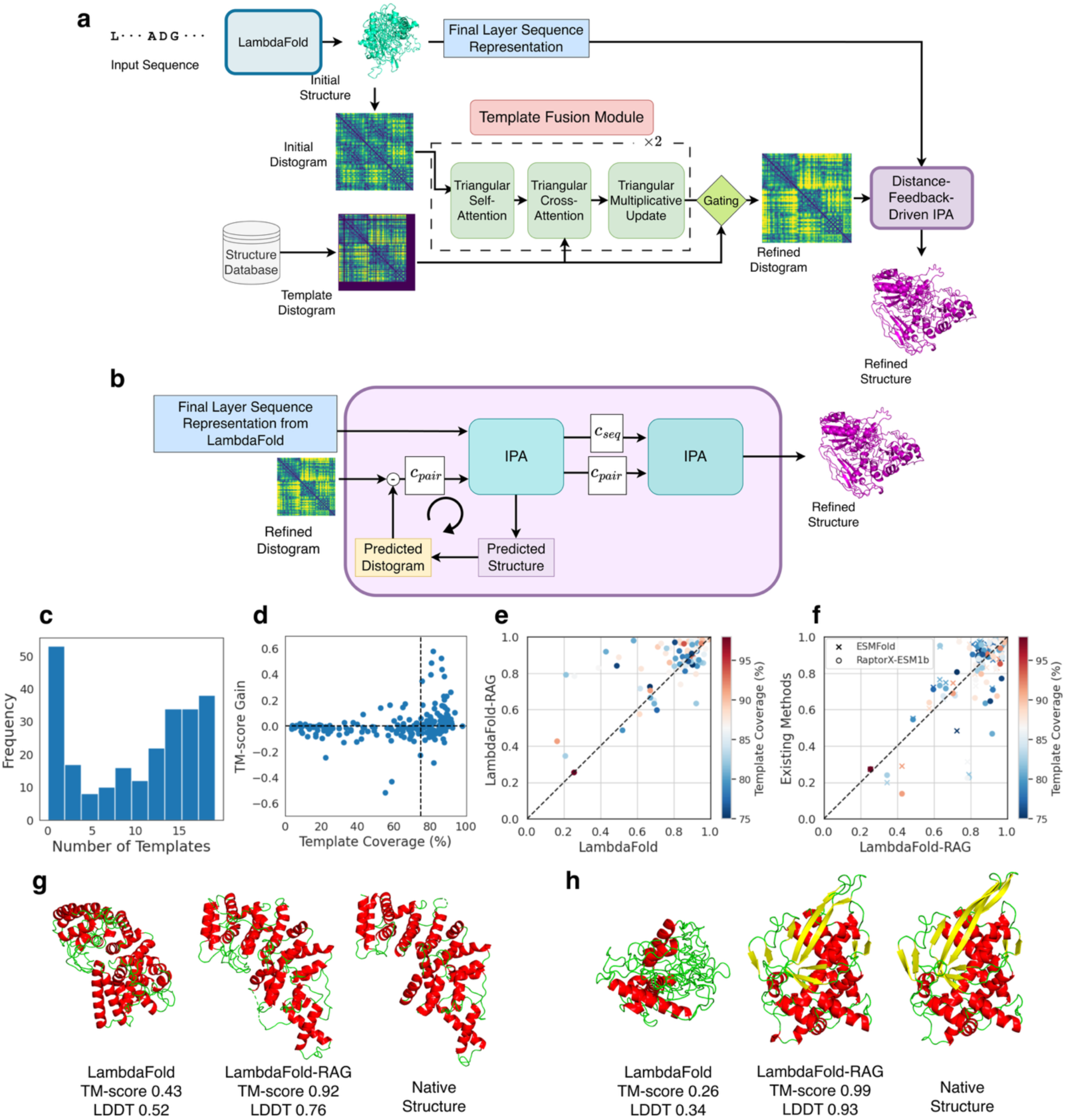
LambdaFold-RAG. **a**. schematic overview of the RAG pipeline. Initial structure predicted by LambdaFold is converted into a distogram and fused with a template distogram retrieved from external databases via the Template Fusion Module. The Template Fusion Module utilizes triangular self-attention, triangular cross-attention (incorporating template data), and triangular multiplicative updates, followed by a gating mechanism to produce a refined distogram. The resulting refined distogram guides a Distance-Feedback-Driven Invariant Point Attention (IPA) module to produce the final refined structure. **b**. detailed view of the Distance-Feedback-Driven IPA. This module iteratively minimizes the error between the current predicted distogram and the template distogram refined by Template Fusion Module, followed by a final IPA layer to refine full atomic geometry. **c**. the number of templates identified for the 244 combined CASP14 and CAMEO benchmark proteins. 231 had templates available. **d**. the change of TM-score of structure models relative to the template coverage. **e**. Performance comparison (TM-score) between the base LambdaFold and the LambdaFold-RAG version, which uses templates. 90 targets that had a template with at least 75% sequence coverage were plotted. The color gradient indicates template coverage percentage. The mean TM-score improved from 0.80 to 0.85 by using templates with the RAG pipeline. **f**. TM-score comparison between LambdaFold-RAG with ESMFold and RaptorX-ESM1b on the 90 targets. The average: LambdaFold-RAG, 0.85; ESMFold, 0.86, RaptorX-ESM1b, 0.83. **g**. an example of modeling difference between LambdaFold and LambdaFold-RAG. The target shown is 7fiw_B. A baseline LambdFold model and the corresponding LambdaFold-RAG, and the native structures are shown. TM-score and LDDT are shown below the models. **h**. Modeling for T1037 with LambdaFold and LambdaFold-RAG.

The template fusion module consists of two fusion blocks, each comprising three triangular layers (**Fig. 5a**). Each block first applies triangular self-attention to the LambdaFold-predicted distogram, followed by triangular cross-attention with the template distogram while masking missing template entries. A triangular update and gating mechanism then selectively incorporates available template information while retaining LambdaFold predictions where template information is absent. The resulting fused distogram therefore combines the predicted and template-derived distance information for subsequent structure refinement (see Methods).

**Fig. 5b** illustrates the distance-feedback-driven IPA module. At each iteration, a distogram is computed from the current predicted structure (predicted distogram in the figure) and compared with the refined template distogram to obtain distance errors. The sequence representation is inherited from LambdaFold (blue box), while the pair representation is constructed from these distance errors. The module was trained using a Cβ displacement loss to enforce the target distance constraints. Because this optimization may distort local protein geometry, a subsequent IPA layer is trained with a standard AF2-style loss to refine the full atomic structure and restore structural integrity (Methods).

Of the 244 target proteins in the combined CASP14 and CAMEO datasets, templates were identified for 199 using the template search protocol with strict temporal cutoffs (May 18, 2020 for CASP14 and April 2, 2022 for CAMEO) to ensure that only structures available at the time of each benchmark were included (Fig. 5c). On average, 11.56 templates were retrieved per target. However, the identified templates include that cover only part of the query sequence, limiting the structural information they provide. **Fig. 5d** shows the change in TM-score with template coverage, defined as the percentage of target residues covered by structural information from the combined templates. High-coverage templates tend to provide sufficient distance information to guide predictions toward the correct fold, whereas low-coverage templates may introduce uncertainty and occasionally degrade an otherwise accurate prediction. **Fig. 5d** shows that the majority of targets with >75% template coverage improved TM-scores with LambdaFold-RAG (individual data in **Supplementary Table S14**).

In **Fig. 5e**, we compared LambdaFold and LambdaFold-RAG on 90 targets with >75% template coverage. LambdaFold-RAG improved the TM-score by more than 0.01 for 56 targets (62.2%), increasing the average TM-score from 0.80 to 0.85. Notably, several challenging targets with initially incorrect folds (TM-score <0.5) were rescued to the correct topology, including largest improvements observed in T1050 (0.39 to 0.83), 7fiw_B (0.43 to 0.92), 7eqs_A (0.21 to 0.79), 7ms2_A (0.49 to 0.84), and 7n45_A (0.25 to 0.78). These results demonstrate that high-coverage templates can provide effective structural guidance and substantially improve LambdaFold predictions, particularly for difficult targets. When compared with ESMFold and RaptorX-ESM1b models on these 90 targets (**Fig. 5f**), LambdaFold-RAG achieves a higher average TM-score than RaptorX-ESM1b (0.85 vs. 0.83) and performs almost on par with ESMFold (0.85 vs 0.86) (individual data in **Supplementary Table S15**).

**Fig. 5g and 5h** show two examples illustrating the substantial improvement achieved by LambdaFold-RAG. For the α-helical protein 7fiw_B (**Fig. 5g**), LambdaFold correctly folded only approximately half of the structure (TM-score: 0.43), whereas template guidance recovered the correct overall topology, increasing the TM-score to 0.92. We observe the effect of using a native distogram as a template in the second example, an α/β protein (**Fig. 5h**), LambdaFold correctly modeled only part of the α-helical region, while the β-sheet was largely misfolded (TM-score: 0.26). LambdaFold-RAG dramatically improved the prediction, recovering nearly the entire structure with a TM-score of 0.99. The distance-feedback IPA (**Fig. 5b**) effectively follows the distogram prompt. In standard AF2 IPA, the pair representation enters only as a bias term, limiting its influence relative to the sequence representation; consequently, even ground-truth templates provide only modest improvements [61]. In contrast, our iterative, error-driven integration gives template-derived pair information a more direct and dominant role in structure refinement.

## Discussion

We presented Prot-LAMBDA, a PLM enhanced with explicit inter-residue distance awareness through a pretraining strategy that aligns embedding differences with spatial distances. Although PLMs have achieved remarkable success in many bioinformatics applications, including protein structure prediction [20], they are trained primarily by masked residue prediction and therefore encode structural information only implicitly through sequence context. In this work, we aimed to directly incorporate inter-residue distance constraints into the representation learning process, which is critical to reproduce protein structures.

As intended, Prot-LAMBDA consistently outperformed ESM despite using approximately one-fifth as many parameters across a range of structure-related prediction tasks, including secondary structure, residue contacts, dihedral angles, solvent accessibility, and protein fold classification. Furthermore, when coupled with the same structure prediction module and trained under identical conditions, Prot-LAMBDA achieved higher tertiary structure prediction accuracy than ESM (**Fig. 2**). These results demonstrate that Prot-LAMBDA learns substantially richer structural representations than ESM.

LambdaFold, which we built with Prot-LAMBDA, did not exceed the performance over ESMFold; however, it showed on par performance in challenging protein targets and targets that are strictly non-redundant from the training set. Importantly, the results also suggest that ESMFold memorizes structures that were used in training and apparent good performance are coming at least partially from memorization.

Current limitation of LambdaFold is its relatively shallow folding trunk, which limits recycling and results in only modest performance gains. This design was primarily driven by the computational resources available during model development. Future work will investigate deeper trunk architectures, scale both Prot-LAMBDA and LambdaFold with larger training datasets, and expand the retrieval database with additional AFDB templates, as demonstrated by the improved performance of LambdaFold-RAG. We also plan to extend the framework to multimeric complexes and other biomolecules.

## Acknowledgements

This work was partly supported by the National Institutes of Health (R35GM158267, R21AI187928) and the National Science Foundation (NSF) (IIS-2211598, DMS-2151678, DBI-2146026, and DBI-2422620) to DK; the National Institutes of Health (R35GM158094, R01GM134020) and the National Science Foundation (NSF) (DBI-2238093, DBI-2422619, IIS-2211597, and MCB-2205148) to MX; and the Research Support Project for Life Science and Drug Discovery (Basis for Supporting Innovative Drug Discovery and Life Science Research BINDS) from AMED under Grant Number JP25ama121028 to KT. This work used Delta GPU cluster through allocation BIO240337 from the Advanced Cyberinfrastructure Coordination Ecosystem: Services & Support (ACCESS) program, which is supported by NSF (OAC2138259, OAC2138286, OAC2138307, OAC2137603, and OAC2138296).

## Code Availability

The Prot-LAMBDA code along with pretrained weights are freely available for academic use from GitHub at https://github.com/kiharalab/Prot-LAMBDA. The trained model weights are made available at https://huggingface.co/KiharaLab/ProtLAMBDA.

## Data Availability

Raw data for the figures are provided in Tables S4-S15 in a separate Excel file. Predicted structures, contact maps and distograms are available at Zenodo (10.5281/zenodo.21984993).

## Author Contributions

D.K. conceived the study. N.I. designed and implemented algorithms, conducted the experiments, and analyzed the data. Z.Z. contributed to the implementation of the AF2-based IPA and FAPE loss code. Y.K. and N.I. performed dataset preparation and processing. N.I. drafted the manuscript; D.K. thoroughly edited the manuscript. K.T. and M.X. participated in model training process, discussion, and editing. All authors read and approved the final manuscript.

## Conflict of Interests

The authors declare no conflict of interests.

## Methods

### Training dataset

We constructed a mixed training dataset comprising both experimentally determined structures from the Protein Data Bank (PDB) and computationally predicted structures from the AlphaFold Database (AFDB). As CASP14 and CAMEO serve as our primary evaluation benchmarks, we carefully designed the training set to ensure non-redundancy with respect to these test datasets.

For the PDB-derived dataset, we included all protein chains released before January 2020 with a resolution ≤9 Å and sequence length >20 residues, following the protocol of ESMFold [28]. We removed redundant sequences with ≥20% sequence identity, ignored hetero atoms and ligands, and, unlike previous studies, included NMR structures by selecting the first model from each ensemble. For proteins with identical sequences, only one chain was retained. After excluding chains sharing >25% sequence identity with proteins in the CASP14 or CAMEO test sets, the dataset contained 98,908 chains (**Supplementary Table S1**). These were subsequently clustered at 40% sequence identity using MMseqs2, yielding 32,137 clusters for efficient training.

For the AFDB-derived dataset, we sampled sequences from UniClust30 (release 2018_08) [62]. Clusters were sorted in descending order by size, and we observed that the mean pLDDT decreased as cluster size increased. We therefore retained only clusters containing more than 30 sequences and selected one representative sequence from each cluster. Following ESMFold, we excluded proteins with a mean pLDDT <70 and masked residues with per-residue pLDDT <70. Proteins sharing more than 25% sequence identity with CASP14 or CAMEO targets were also removed. The resulting dataset contains 310,267 proteins (**Supplementary Table S2**). No further clustering was performed because the sequences were already clustered in UniClust30.

To the best of our knowledge, our method is the only one that explicitly enforces this level of non-redundancy between the training data and benchmark test sets. Although this places our model at a relative disadvantage compared with previous approaches, it provides a more rigorous assessment of generalization. Furthermore, unlike methods such as ESMFold, which are trained on up to ∼14 million proteins, we intentionally use a substantially smaller training set to emphasize representation learning and architectural inductive bias rather than data memorization, thereby better isolating the model’s intrinsic generalization capability.

We used the same validation set as ESMFold, consisting of 776 CAMEO proteins released between August 2021 and January 2022 (**Supplementary Table S3**).

### Casp14 and CAMEO test sets

Following prior works [15, 28], we evaluated our method on a benchmark comprising 50 CASP14 proteins (the target T1044, with 2180 residues was excluded since it results in out of memory even by the ESM language models) and 194 proteins from the CAMEO dataset released between April 1 and June 25, 2022. For evaluations of protein language model representations, including contact map prediction and AF2 IPA-based structure prediction, we used the full benchmark set.

For evaluating distance prediction, we focused on 37 FM and FM/TBM domains in the CASP14 test set, as defined in previous works [53, 54]. The domains considered are: T1027-D1, T1029-D1, T1031-D1, T1033-D1, T1035-D1, T1037-D1, T1038-D1, T1038-D2, T1039-D1, T1040-D1, T1041-D1, T1042-D1, T1043-D1, T1046s1-D1, T1047s1-D1, T1047s2-D1, T1047s2-D3, T1049-D1, T1052-D3, T1053-D1, T1053-D2, T1055-D1, T1058-D1, T1061-D1, T1061-D2, T1064-D1, T1065s2-D1, T1070-D1, T1074-D1, T1080-D1, T1082-D1, T1090-D1, T1093-D1, T1093-D3, T1094-D2, T1096-D1, T1096-D2

However, because a substantial fraction of the benchmark proteins shares sequence similarity with the ESMFold training dataset, we restricted our 3D structure prediction evaluation to a non-redundant subset to enable a more rigorous assessment of generalization. Specifically, we clustered the CASP14 and CAMEO proteins at a 25% sequence identity threshold using MMseqs2, resulting in a benchmark of 30 non-redundant proteins. The resulting set comprises:

CASP14 (12 proteins): T1030, T1031, T1033, T1035, T1037, T1038, T1039, T1040, T1042, T1043, T1049, T1055

CAMEO (18 proteins): 7zro_A, 7mnk_A, 7mo1_B, 7mo3_B, 7r24_A, 7mq4_A, 7rkc_A, 7sxb_A, 7opb_D, 7mla_B, 7qao_A, 7qap_A, 7tni_C, 7n3t_C, 7pno_D, 7t7y_A, 7eqb_A, 7wwr_A

### Prot-LAMBDA architecture

We adopted the BERT-style ESM2 architecture as the backbone protein language model (**Fig. 1a**). A pretrained ESM2-650M model (33 transformer layers, embedding dimension 1280), trained on UniRef50 (version 2021_04), was used for initialization. During distance-aware pretraining, only the top 33%, i.e., 11 transformer layers were fine-tuned, while the remaining layers were frozen.

Residue-level pair representations were constructed from the element-wise absolute difference between residue embeddings. The pair representation was projected from 1280 to 512 dimensions and refined using a single axial attention layer (8 attention heads), in which row-wise and column-wise attention outputs were summed. Rotary positional embeddings (RoPE) were applied along both axes, and dropout (p=0.05) was used for regularization. Binary residue contact probabilities were predicted from the final pair representation using a single linear projection implemented as a 1 × 1 convolution.

### Prot-LAMBDA loss function

The Prot-LAMBDA model was trained jointly using masked language modeling (MLM) and contact map prediction objectives. The MLM loss was computed as the cross-entropy over masked residues following the standard BERT training protocol. Contact map prediction was optimized using binary cross-entropy. The final training objective was the sum of the two losses.

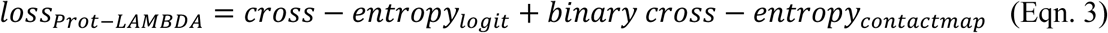

### Prot-LAMBDA training

To mitigate catastrophic forgetting during joint optimization, we adopted a curriculum learning strategy for contact map supervision. The masked language modeling objective followed the standard protocol, with 15% of residues randomly masked throughout training. For contact prediction, the loss was initially computed on a random 25% subset of residue pairs, which was linearly increased to 50% over the first 20 epochs and to 100% over epochs 20-40. Full contact map supervision was then applied for the remaining 40 epochs.

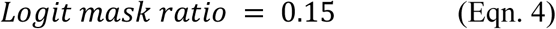

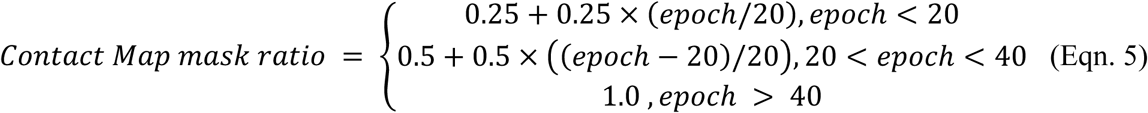

The model was trained for 80 epochs using the AdamW optimizer with an effective batch size of 1,024. The learning rate was annealed from 10^−3^ to 10^−6^ using a cosine schedule. The best model was selected based on validation performance. Training was performed using fixed-length sequence crops of 256 residues. We found that the RoPE embedding could generalize effectively beyond this length and therefore did not train on longer sequence crops.

### Prot-LAMBDA downstream applications

For downstream structure feature prediction (Fig. 2e), residue-level embeddings were used as input to a linear probe, except for fold prediction, where global mean-pooled embeddings were used. A linear layer was trained using cross-entropy for classification tasks and mean squared error for regression tasks. Optimization was performed using Adam for 10 epochs with a learning rate of 10^-3^.

### PLM-only 3D structure prediction

For PLM-only de novo structure prediction (**Fig. 2f, 2g**), frozen PLM embeddings and attention maps were projected to the input dimensions of the AF2 structure module and used as sequence and pair representations, respectively (**Supplementary Fig. 2**). Residue embeddings from Prot-LAMBDA and the ESM2-650M and ESM2-3B models were projected to 384-dimensional sequence representations, whereas attention maps (axial attention for Prot-LAMBDA and self-attention for ESM2) were projected to 128-dimensional pair representations.

The projection layers were randomly initialized, whereas the AF2 structure module was initialized from pretrained weights. During training, only the projection layers and the structure module were updated, while the PLMs remained frozen.

All models were trained on the same set of 98,908 PDB structures for 30 epochs using the standard AF2 loss, excluding the violation loss.

### DistFormer architecture

Our DistFormer architecture is based on the Evoformer or folding trunk used in AF2-style structure predictors (**Fig. 3a**). Sequence representations were updated using pair-biased self-attention, followed by pair representation updates and triangular attention to maintain geometric consistency.

To enhance pair representations, all-pairs sequence features were processed using regional and sparse attention before integration with the pair representation. Regional attention operated on non-overlapping m×m windows (m=8) to capture local interactions, whereas sparse attention attended over regularly sampled residue grids to capture long-range interactions. The outputs of the two attention modules were concatenated and fused with the pair representation. Each DistFormer block contained two regional-sparse attention modules while retaining the standard sequence attention and triangular attention operations of the Evoformer.

DistFormer comprised 6 layers organized into three sequential stages of 2 layers each. The three stages predicted distograms over the range 2.3125-21.6875 Å using 4, 16, and 64 distance bins, respectively. A shared IPA-based structure module generated 3D structures from the sequence and pair representations at the end of each stage. To account for the increasing complexity, sequence and pair representation dimensions were scaled across stages from (512, 192) to (640, 224) and (768, 256), respectively.

### DistFormer loss-function

We adopted a loss formulation closely aligned with that used in AF2. The overall loss is defined as:

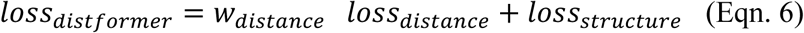

Here, *loss_structure_* corresponds to the standard AlphaFold structural loss, for which we use the same components as implemented in OpenFold [63]. The only modification is that we include the violation loss throughout training, with a reduced weight of 0.25.

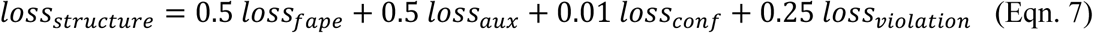

The *loss_structure_* is designed to explicitly account for both probabilistic accuracy and distance error magnitude. Unlike standard cross-entropy loss, which considers only the predicted distribution, our formulation incorporates a multiplicative combination of Huber loss and cross-entropy. The Huber loss provides a balanced alternative to mean squared error (MSE) and mean absolute error (MAE): for errors below a threshold δ, it behaves like MSE, while for larger errors it transitions to a MAE penalty, thereby reducing the influence of outliers and stabilizing training. Our formulation encourages the model to assign high probability to the correct distance bin while simultaneously penalizing large deviations in predicted distances:

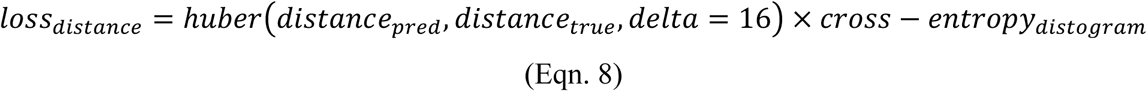

Finally, *w_distance_* is a scaling factor applied to the distance loss. Due to the inclusion of the Huber term, the magnitude of *loss_structure_* increases in later Distformer stages, potentially overwhelming the structural loss. To maintain stable training, we apply stage-dependent weights of 1.0, 0.5, and 0.1 for stages 1, 2, and 3, respectively.

### DistFormer training

DistFormer was trained on the Prot-LAMBDA training dataset using the loss described above (Eq. 8). The three stages were trained sequentially. The first stage and a shared IPA-based structure module were trained using features extracted from the frozen Prot-LAMBDA model. After convergence, the first stage was frozen and the second stage was trained, followed by the third stage using the same procedure. The IPA-based structure module was shared across all three stages.

Each stage was trained for 30 epochs using the AdamW optimizer with an effective batch size of 1,024. The learning rate was annealed from 10−3 to 10−6 using a cosine schedule. The best model was selected based on validation performance.

### Fisher Discriminant Ratio (FDR)

Fisher discriminant ratio (FDR) provides a quantitative measure of class separability by comparing the separation of class centroids with the variability of samples within each class. For a dataset comprising *C* classes, let class *i* contain *n_i_* samples, and let µ_i_ and σ_i_ denote the mean and standard deviation of a given feature within class *i*, respectively. Let μ denote the global mean across all classes. The FDR of the feature is defined as

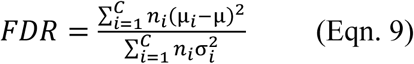

The numerator quantifies between-class variability, measuring how far the mean of each class is from the global mean, with each class weighted by its number of samples. The denominator quantifies within-class variability, measuring the spread of feature values among samples belonging to the same class. Thus, FDR is high when class means are well separated while samples within each class remain relatively compact, indicating that the feature provides strong discriminatory information. Conversely, a low FDR indicates that the class means are relatively close compared with the variability within classes, resulting in greater overlap between classes.

### Structural template generation

Template retrieval was performed using the standard AF2 template search pipeline implemented in ColabFold [60]. For each query sequence, experimentally determined structures were retrieved from the PDB70 database using the MMseqs2-based search pipeline [58], and the top 20 candidate templates were refined using HHsearch [59]. Accepted templates were converted into the standard AF2 template features, from which the template distogram was extracted. The distogram encodes pairwise Cβ distances (Cα for glycine) using 39 distance bins spanning 3.25-50.75 Å with an additional overflow bin.

Only the template distogram was used as input to the retrieval-augmented generation (RAG) module. During evaluation, template retrieval was restricted to PDB entries released before the corresponding target release date to prevent data leakage.

### Template fusion module

The Template Fusion Module (**Fig. 5a**) consists of two sequential refinement blocks. Each block applies triangular self-attention, followed by triangular cross-attention in which the predicted distogram serves as the query and the template distogram as the key and value. Missing template entries were excluded using attention masks. The refined representation was further updated using a triangular multiplicative update. After the two refinement blocks, a gating operation selected the template distogram for residue pairs with available template information and the predicted distogram otherwise, producing the final fused distogram.

The Template Fusion Module was trained using a simulated distogram inpainting task. During training, 5-15% of template distogram entries were randomly masked, and the model was optimized to reconstruct the masked distances using the distogram cross-entropy loss. The module was trained for 30 epochs on the 98,908 PDB structures in the training set.

### Distance-feedback-driven IPA

The Distance-Feedback-Driven IPA module (**Fig. 5b**) imposes the refined template distogram into the initially predicted 3D protein structure. The module consists of two sequential IPA blocks. Given the sequence representation *c_seq_* from LambdaFold and the refined template distogram D_ref_, the first IPA block iteratively updates the Cβ coordinates (Cα for glycine). At iteration *t*, a predicted distogram is computed from the current coordinates.

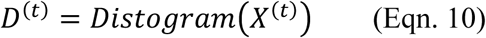

Where *X*^(*t*)^ denotes the current predicted Cβ coordinates. The pair representation is then defined as the discrepancy between the refined template distogram and the current prediction,

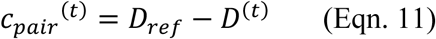

which provides explicit geometric feedback indicating how each residue pair should be corrected. The IPA block updates the coordinates according to

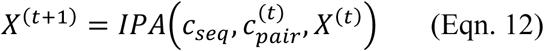

Where the sequence representation *c_seq_* remains fixed throughout the refinement process. By repeatedly recomputing the distogram from the updated coordinates and feeding back the residual distance error, the predicted structure progressively satisfies the template-guided distance constraints.

The refined Cβ coordinates were subsequently passed to the second IPA block, which predicted the full-atom structure using the standard AF2 formulation.

The two IPA blocks were trained sequentially. The first block was optimized using a mean squared error loss on the predicted Cβ coordinates, whereas the second block was trained using the standard AF2 structural loss. Training was performed for 30 epochs on the 98,908 PDB structures in the training dataset.

### Computational resource

Prot-LAMBDA was trained on four NVIDIA A100 GPUs (40 GB memory each) for approximately 2000 GPU-hours. Early ablation studies and architecture development for LambdaFold were performed on eight NVIDIA V100 GPUs (16 GB memory each). The final LambdaFold and LambdaFold-RAG models were trained on four NVIDIA A100 GPUs (40 GB memory each) for approximately 7500 GPU-hours.

## Notes

### Competing Interest Statement

The authors have declared no competing interest.

