## Supplementary Figure for "Prot-LAMBDA: Explicit Distance Learning Enhances Structural Reasoning in Protein Language Models"

**Affiliations**

**Supplementary Figure 1:** Examples of predicted protein contact maps. These correspond to the same proteins shown in Fig. 1D–G, with the addition of ESM2-650M model predictions.

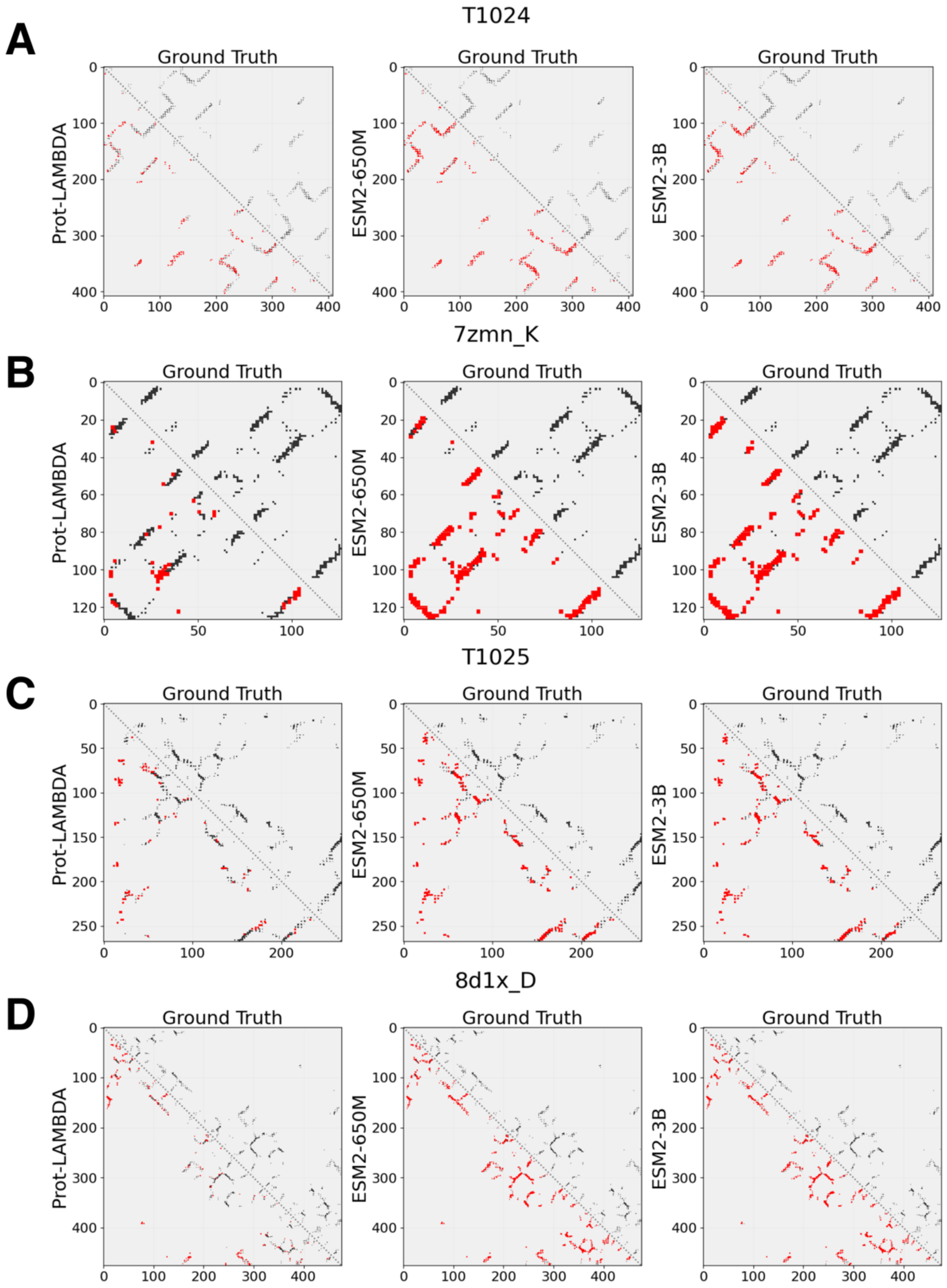

A) **CASP14 target T1024 (LmrP, PDB ID: 6t1z_A):** An α-class protein, length: 408 amino acids (aa).

**B) RECQL5 helicase (PDB ID: 7zmn_K):** A β-class protein, length: 127 aa.

**C) T1025 (AtmM, PDB ID: 6uv6_A):** An α/β-class protein, length: 268 aa.

**D) Aminopeptidase A (PDB ID: 8d1x_D):** An α/β-class protein, length: 476 aa.

Accuracy metrics of these targets are summarized below:

| Protein Target | Metric | Short | | | Medium | | | Long | | |
|---|---|---|---|---|---|---|---|---|---|---|
| | | **Prot-LAMBDA** | **ESM2-650M** | **ESM2-3B** | **Prot-LAMBDA** | **ESM2-650M** | **ESM2-3B** | **Prot-LAMBDA** | **ESM2-650M** | **ESM2-3B** |
| **T1024** | Precision @L | 0.76 | 0.67 | 0.73 | 0.87 | 0.7 | 0.7 | 0.74 | 0.67 | 0.69 |
| | Recall | 0.78 | 0.58 | 0.73 | 0.88 | 0.66 | 0.71 | 0.69 | 0.36 | 0.41 |
| | F1 Score | 0.71 | 0.6 | 0.71 | 0.87 | 0.7 | 0.7 | 0.69 | 0.48 | 0.54 |
| **7zmn_K** | Precision @L | 0.9 | 0.67 | 0.57 | 0.87 | 0.67 | 0.61 | 0.91 | 0.65 | 0.54 |
| | Recall | 0.84 | 0.41 | 0.35 | 0.86 | 0.48 | 0.4 | 0.8 | 0.26 | 0.2 |
| | F1 Score | 0.89 | 0.55 | 0.5 | 0.87 | 0.62 | 0.53 | 0.82 | 0.38 | 0.32 |
| **T1025** | Precision @L | 0.9 | 0.67 | 0.75 | 0.83 | 0.63 | 0.65 | 0.97 | 0.8 | 0.79 |
| | Recall | 0.9 | 0.65 | 0.75 | 0.91 | 0.5 | 0.56 | 0.77 | 0.42 | 0.45 |
| | F1 Score | 0.9 | 0.72 | 0.78 | 0.82 | 0.59 | 0.64 | 0.78 | 0.55 | 0.57 |
| **8d1x_D** | Precision @L | 0.81 | 0.52 | 0.55 | 0.83 | 0.42 | 0.49 | 0.99 | 0.7 | 0.71 |
| | Recall | 0.77 | 0.31 | 0.36 | 0.78 | 0.26 | 0.28 | 0.81 | 0.24 | 0.27 |
| | F1 Score | 0.82 | 0.42 | 0.47 | 0.82 | 0.38 | 0.41 | 0.83 | 0.37 | 0.4 |

Here, we report the precision@L (precision computed using the top L predicted contact probabilities, where L is the length of the protein), recall, and F1 score across short-, medium-, and long-range contacts. These ranges correspond to contacts between residues with a sequence distance of 6–12, 12–24, and greater than 24, respectively.

**Supplementary Figure 2:** Overview of the de novo protein structure prediction architecture used to evaluate Prot-LAMBDA embeddings

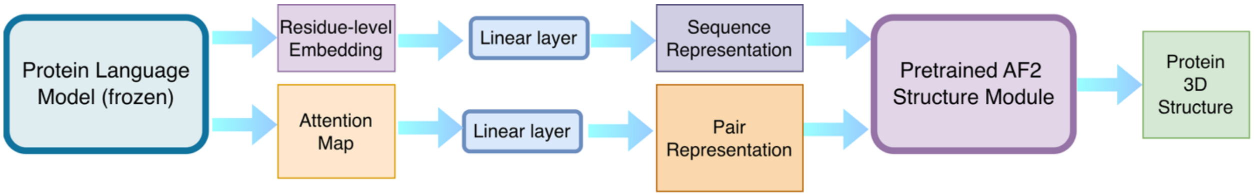


Input protein sequences are first processed by a frozen Prot-LAMBDA model (or frozen ESM2 baselines) to extract high-dimensional residue-level embeddings. Attention maps are also extracted from the protein language models (axial attention for Prot-LAMBDA and standard self-attention for ESM2). To bridge the dimensional mismatch between these representations and the AlphaFold2 structure module, the embeddings and attention maps are projected through separate trainable linear layers to generate the required sequence and pair representations, respectively. The model is initialized with a pretrained AF2 structure module and fine-tuned on 98,908 experimentally determined PDB structures for 30 epochs using the standard AF2 loss functions, excluding the violation loss. Throughout training, the protein language models remain frozen.

**Supplementary Table 1:** Contact Map Performance Assessment on the CASP14 benchmark (corresponding to Fig. 2A). The dataset includes 50 targets. Here we list the short, medium, long-range Precision (Pr), Recall (Re) and F1 scores and present a comparison of Prot-LAMBDA against baseline ESM2-650M and ESM2-3B models. The best scores are highlighted in bold.

| Model | Short Pr | Short Re | Short F1 | Med Pr | Med Re | Med F1 | Long Pr | Long Re | Long F1 |
|---|---|---|---|---|---|---|---|---|---|
| **Prot-LAMBDA** | **0.65** | **0.51** | **0.52** | **0.58** | **0.48** | **0.49** | **0.49** | **0.39** | **0.38** |
| **ESM2-650M** | 0.52 | 0.35 | 0.39 | 0.46 | 0.30 | 0.33 | 0.50 | 0.17 | 0.21 |
| **ESM-3B** | 0.55 | 0.39 | 0.43 | 0.50 | 0.33 | 0.37 | 0.49 | 0.19 | 0.23 |

**Supplementary Table 2:** Contact Map Performance Assessment on CAMEO benchmark (corresponding to Fig. 2B). The dataset includes 194 targets. Here we list the short, medium, long-range Precision (Pr), Recall (Re) and F1 scores and present a comparison of Prot-LAMBDA against baseline ESM2-650M and ESM2-3B models. The best scores are highlighted in bold.

| Model | Short Pr | Short Re | Short F1 | Med Pr | Med Re | Med F1 | Long Pr | Long Re | Long F1 |
|---|---|---|---|---|---|---|---|---|---|
| **Prot-LAMBDA** | **0.75** | **0.70** | **0.71** | **0.76** | **0.71** | **0.72** | **0.67** | **0.60** | **0.62** |
| **ESM2-650M** | 0.65 | 0.42 | 0.50 | 0.65 | 0.43 | 0.50 | 0.65 | 0.24 | 0.33 |
| **ESM-3B** | 0.67 | 0.44 | 0.52 | 0.66 | 0.44 | 0.51 | 0.67 | 0.27 | 0.36 |

**Supplementary Table 3:** Contact Map Performance Assessment on TAPE benchmark (corresponding to Fig. 2C). The dataset includes 40 targets. Here we list the short, medium, long-range precision scores at different level of top hits (L/k, k=1,2,5) and present a comparison of Prot-LAMBDA against existing structurally informed PLMs: Amplify, S-PLM, ESM-2S, SaProt, and ISM, along with the baseline ESM2-650M. The best scores are highlighted in bold.

| Model | Short P@L | Short P@L/2 | Short P@L/5 | Med P@L | Med P@L/2 | Med P@L/5 | Long P@L | Long P@L/2 | Long P@L/5 |
|---|---|---|---|---|---|---|---|---|---|
| **Prot-LAMBDA** | **0.65** | **0.65** | **0.71** | **0.61** | **0.62** | **0.70** | **0.56** | **0.66** | **0.75** |
| **ISM** | 0.62 | 0.62 | 0.67 | 0.60 | 0.61 | 0.68 | 0.49 | 0.57 | 0.69 |
| **SaProt** | 0.57 | 0.57 | 0.64 | 0.53 | 0.55 | 0.66 | 0.48 | 0.60 | 0.74 |
| **S-PLM** | 0.49 | 0.49 | 0.55 | 0.48 | 0.49 | 0.57 | 0.36 | 0.43 | 0.54 |
| **Amplify** | 0.38 | 0.38 | 0.41 | 0.36 | 0.35 | 0.40 | 0.23 | 0.28 | 0.35 |
| **ESM-2S** | 0.46 | 0.46 | 0.50 | 0.46 | 0.47 | 0.54 | 0.36 | 0.43 | 0.52 |
| **ESM2-650M** | 0.45 | 0.45 | 0.50 | 0.45 | 0.45 | 0.54 | 0.35 | 0.42 | 0.52 |

**Supplementary Table 4:** Contact Map Performance Assessment on 15 CASP14-FM targets (corresponding to Fig. 2D). Here we list the medium, long-range precision scores at different level of top hits (L/k, k=1,2,5,10) and present a comparison of Prot-LAMBDA against dedicated single-sequence-based methods such as SPOT-Contact-LM, SSCPred, along with the baseline ESM-1b and profile-based SPOT-Contact. The best scores are highlighted in bold.

| Model | Med P@L | Med P@L/2 | Med P@L/5 | Med P@L/10 | Long P@L | Long P@L/2 | Long P@L/5 | Long P@L/10 |
|---|---|---|---|---|---|---|---|---|
| **Prot-LAMBDA** | **0.25** | **0.26** | 0.29 | 0.33 | 0.15 | **0.19.** | **0.30** | **0.35** |
| **ESM-1b** | 0.11 | 0.16 | 0.20 | 0.23 | 0.07 | 0.09 | 0.12 | 0.17 |
| **SSCpred** | 0.13 | 0.17 | 0.25 | 0.26 | 0.08 | 0.08 | 0.09 | 0.10 |
| **SPOT-Contact-LM** | 0.14 | 0.18 | 0.25 | 0.30 | 0.12 | 0.15 | 0.19 | 0.19 |
| **SPOT-Contact (profile)** | 0.17 | **0.26** | **0.36** | **0.41** | **0.16** | **0.19** | 0.21 | 0.25 |

**Supplementary Table 5:** Performance assessment on downstream structural characterization (corresponding to Fig. 2E). Tasks include prediction of 3-state (SS3) and 8-state (SS8) protein secondary structure, disordered region, PHI and PSI dihedral angles, relative solvent accessibility (RSA), and protein fold topology from the PEER benchmark and comparison against ESM2 models. Metrics include accuracy (ACC ↑) for 3-state (SS3) and 8-state (SS8) secondary structure and protein fold prediction; False Negative Rate (FNR ↓) for disorder; Mean Absolute Error (MAE ↓) for PHI and PSI backbone dihedral angles; and Pearson Correlation Coefficient (PearsonR ↑) for Relative Solvent Accessibility (RSA). Arrows indicate the direction of improved performance.

| **Model** | **SS3 (ACC ↑)** | **SS8 (ACC ↑)** | **Disorder (FNR ↓)** | **PHI (MAE ↓)** | **PSI (MAE ↓)** | **RSA (PearsonR ↑)** | **Fold (ACC ↑)** |
|---|---|---|---|---|---|---|---|
| **ESM2-650M** | 0.8419 | 0.7195 | 0.0209 | 29.8751 | 46.2229 | 0.7131 | 0.3050 |
| **ESM2-3B** | 0.8459 | 0.7228 | 0.0181 | 29.6526 | 45.7025 | 0.7271 | 0.3134 |
| **Prot-LAMBDA** | **0.8551** | **0.7386** | **0.0151** | **28.4337** | **40.9933** | **0.7337** | **0.3203** |

**Table captions for the Supplementary Tables provided as an Excel file**

**Supplementary Table S1. The list of PDB chains used in training**. In the Excel file, the table is labeled as "TableS1" in the Excel file.

**Supplementary Table S2. The list of AFDB proteins used in training**. In the Excel file, the table is labeled as "TableS2" in the Excel file.

**Supplementary Table S3. The list of PDB chains used as validation data**. In the Excel file, the table is labeled as "TableS3" in the Excel file.

**Supplementary Table S4. Correlation between embedding difference and spatial distance for Prot-LAMBDA and ESM2 PLMs on the CASP14 and CAMEO benchmark proteins.** Raw data for Fig 1c, labeled as "TableS4" in the Excel file.

**Supplementary Table S5. Contact map prediction metrics for Prot-LAMBDA and ESM2 PLMs on the CASP14 and CAMEO benchmark proteins.** Raw data for Fig 2a and 2b, labeled as "TableS5" in the Excel file. Predicted contact maps are uploaded to Zenodo (10.5281/zenodo.21984993).

**Supplementary Table S6. Protein 3D structure prediction metrics using frozen PLM embeddings coupled with an IPA structure module on the CASP14 and CAMEO benchmark proteins.** Raw data for Fig 2f and 2g, labeled as "TableS6" in the Excel file. Predicted structures are uploaded to Zenodo (10.5281/zenodo.21984993).

**Supplementary Table S7. Contact formation metrics in predicted 3D structures across the 3 Distformer stages for the CASP14 and CAMEO benchmark proteins.** Raw data for Fig 3d and 3e, labeled as "TableS7" in the Excel file. Predicted structures are uploaded to Zenodo (10.5281/zenodo.21984993).

**Supplementary Table S8. Distogram prediction metrics for LambdaFold and ESMFold on 37 FM/TBM domains from the CASP14 benchmark.** Raw data for Fig 3f, labeled as "TableS8" in the Excel file. Predicted distograms are uploaded to Zenodo (10.5281/zenodo.21984993).

**Supplementary Table S9. Distogram prediction metrics for LambdaFold and ESMFold on 30 non-redundant proteins from the CASP14 and CAMEO benchmark.** Raw data for Fig 3g, labeled as "TableS9" in the Excel file. Predicted distograms are uploaded to Zenodo (10.5281/zenodo.21984993).

**Supplementary Table S10. Protein 3D structure prediction metrics across the 3 Distformer stages for the CASP14 and CAMEO benchmark proteins.** Raw data for Fig 4a, labeled as "TableS10" in the Excel file. Predicted structures are uploaded to Zenodo (10.5281/zenodo.21984993).

**Supplementary Table S11. Protein 3D structure prediction metrics for LambdaFold, RaptorX-ESM1b and ESMFold on the CASP14 and CAMEO benchmark proteins.** Raw data for Fig 4b and 4c, labeled as "TableS11" in the Excel file. Predicted structures are uploaded to Zenodo (10.5281/zenodo.21984993).

**Supplementary Table S12. Relation between perplexity and TM-score for LambdaFold and ESMFold on the CASP14 and CAMEO benchmark proteins.** Raw data for Fig 4d, labeled as "TableS12" in the Excel file. Predicted structures are uploaded to Zenodo (10.5281/zenodo.21984993).

**Supplementary Table S13. Protein 3D structure prediction metrics for LambdaFold, RaptorX-ESM1b and ESMFold on the 30 non-redundant proteins from CASP14 and CAMEO benchmark.** Raw data for Fig 4e and 4f, labeled as "TableS13" in the Excel file. Predicted structures are uploaded to Zenodo (10.5281/zenodo.21984993).

**Supplementary Table S14. Template extraction statistics for the proteins from CASP14 and CAMEO benchmark.** Raw data for Fig 5c and 5d, labeled as "TableS14" in the Excel file. Predicted structures are uploaded to Zenodo (10.5281/zenodo.21984993).

**Supplementary Table S15. 3D structure prediction performance evaluation for LambdaFold-RAG.** Raw data for Fig 5e and 5f, labeled as "TableS15" in the Excel file. Predicted structures are uploaded to Zenodo (10.5281/zenodo.21984993).
